# Characterizing the growing microorganisms at species level in 46 anaerobic digesters at Danish wastewater treatment plants: A six-year survey on microbiome structure and key drivers

**DOI:** 10.1101/2020.06.07.138891

**Authors:** Chenjing Jiang, Miriam Peces, Martin H. Andersen, Sergey Kucheryavskiy, Marta Nierychlo, Erika Yashiro, Kasper S. Andersen, Rasmus H. Kirkegaard, Liping Hao, Jan Høgh, Aviaja A. Hansen, Morten S. Dueholm, Per H. Nielsen

## Abstract

Anaerobic digestion (AD) is a key technology at many wastewater treatment plants (WWTPs) for converting surplus activated sludge to methane-rich biogas. However, the limited number of surveys and the lack of comprehensive data sets have hindered a deeper understanding of the characteristics and associations between key variables and the microbiome composition. Here, we present a six-year survey of 46 anaerobic digesters, located at 22 WWTPs in Denmark, which is the largest known study of the microbial ecology of AD at WWTPs at a regional scale. For three types of AD (mesophilic, mesophilic with thermal hydrolysis pretreatment, and thermophilic), we present the typical value range of 12 key parameters including operational variables and performance parameters. The bacterial and archaeal microbiomes were analyzed at species-level resolution using amplicon sequencing in >1,000 samples and the new ecosystem-specific MiDAS 3 reference database. We detected 42 phyla, 1,600 genera and 3,584 species in the bacterial microbiome, where 70% of the genera and 93% of the species represented uncultivated taxa that were only classified based on MiDAS 3 *denovo* placeholder taxonomy. More than 40% of the 100 most abundant bacterial species did not grow in the digesters and were only present due to immigration with the feed sludge. Temperature, ammonium concentration, and pH were the main drivers shaping the microbiome clusters of the three types of ADs for both bacteria and for archaea. Within mesophilic digesters, feed sludge composition and other key parameters (organic loading rate, biogas yield, and ammonium concentration) correlated with the growing bacterial microbiome. Furthermore, correlation analysis revealed the main drivers for specific species among growing bacteria and archaea, and revealed the potential ecological function of many novel taxa. Our study highlights the influence of immigration on bacterial AD microbiome. Subsetting the growing microbes improves the understanding of the diversity and main drivers of microbiome assembly, and elucidates functionality of specific species-level microorganisms. This six-year survey provides a comprehensive insight into microbiome structure at species level, engineering and ecological performance, and a foundation for future studies of the ecological significance/characteristics and function of the novel taxa.

## Introduction

Anaerobic digestion (AD) is successfully employed worldwide to convert organic feedstock into biogas by anaerobic mixed microbial communities. As a key technology at wastewater treatment plants (WWTP), AD is used to reduce and stabilize the primary and waste-activated sludge by generating methane for bioenergy production. Moreover, AD can be used as a platform for the recovery of value-added compounds (e.g., phosphorus, nitrogen, volatile fatty acids) [1,2]. Thus, it is an important step in the development of circular economy at the WWTPs. The conversion of organic feedstock is carried out by the AD microbiome, a complex network of hydrolyzing and fermenting bacteria, specialized acidogenic and acetogenic syntrophs, and methanogenic archaea [3], which is shaped by stochastic (birth-death immigration) and deterministic (microbial competition, operation and environment) factors [4,5]. Hence, a good understanding of the microbial ecology in digesters is essential for informed control and manipulation of the process for optimal performance.

AD harbours a complex microbial network which is ideal for identifying diversity trends in constrained microbial community structures. Research has shown that the operational parameters, including temperature, substrate type, organic loading rate (OLR), and sludge retention time (SRT) are vital factors for determining the microbiome structure [6–12]. Other parameters, such as ammonia concentration and salinity, are also thought to be significant drivers shaping the microbiome [7,13–15]. Additionally, the microorganisms immigrating with the feed sludge should not be overlooked. Most of them do not grow or contribute to the ecological functions in the system, but they still account for a significant fraction of sequencing reads identified by 16S rRNA gene amplicon sequencing [7,9,13].

However, most of these findings are based on investigations across various AD substrate types, such as manure, food waste, and wastewater sludge, where large differences in growth conditions are observed. Whether the same drivers are also important among digesters at WWTPs is unclear. The AD performance can be highly variable between different WWTPs, but how this links to different microbiomes and growth conditions is poorly described for full-scale systems.

The quantitative relationships between specific microorganisms and key parameters in AD can be evaluated by multiple linear regression (MLR). Most studies have focused on linear associations between methanogenic populations (i.e., characterized by the *mcrA* gene) and specific methanogenic activities [17–20]. However, the traditional MLR fails when the number of predictors is comparable to, or larger than, the number of observations, and when there is high collinearity in predictors. Projection-based methods for analysis of multivariate data, such as Partial Least Squares (PLS) regression, stand as promising techniques to evaluate the links between specific microorganisms and key parameters, lessening the shortcomings of traditional methods [21].

To provide insightful links between the AD microbiome and its performance, it is crucial to obtain a high phylogenetic resolution and good taxonomic classification at all ranks. A high phylogenetic resolution can be obtained by using amplicon sequence variants (ASVs) [22,23] instead of operational taxonomic units (OTUs) typically clustered at 97% similarity thresholds, and by using an ecosystem-specific, high-quality 16S rRNA gene reference database for taxonomic classification. We have developed MiDAS 3, a comprehensive ecosystem-specific reference database for activated sludge and anaerobic digesters which provides a taxonomic classification at all ranks for all sequences based on an improved and automated classification system (AutoTax) [24,25]. The MiDAS 3 reference database is based on full-length 16S rRNA gene ASVs (FL-ASVs) obtained from Danish WWTPs and digesters, but can be applied to similar systems worldwide [24]. MiDAS 3 improves the classification of prokaryotic microorganisms found in AD compared to other public reference databases (SILVA [26], Greengenes [27], and RDP [28]), which lack reference sequences for many taxa and high taxonomic resolution, often resulting in poor classification (**Figure S1**). Application of MiDAS 3 for the study of AD communities offers a possibility of finding the link between identity and function of species-level taxa. Species names provide stable taxa identifiers independent of the data set, thus, allowing cross-study comparisons.

The aims of our study are threefold. Firstly, we describe the typical operational parameters and performance values of three different types of AD at WWTPs (i.e., mesophilic AD, mesophilic AD with pre-treatment (thermal hydrolysis) of waste activated sludge, and thermophilic AD). Secondly, we present the microbial communities in the AD systems (with focus on the growing microbes), for the first time at species level, and make this publicly available on the MiDAS website (https://www.midasfieldguide.org/guide). And thirdly, by focusing on species-level microbiome, we analyse the correlations between key AD parameters and microbiome structure in mesophilic digesters, which are the most common digesters in Denmark at WWTPs.

## Methods

### Anaerobic digesters and sample collection

The survey was conducted during the period 2011 – 2016 in 46 anaerobic digesters at 22 WWTPs across Denmark, which were operated under mesophilic (MAD), mesophilic with thermal hydrolysis pretreatment of feedstock (THP-MAD), or thermophilic (TAD) conditions (see **Table S1** for information of digesters). During the six years of survey, all plants reported minor fluctuations in substrate amounts and composition, but no major changes of operating conditions were introduced, except for Aaby and Aalborg East, which switched from mesophilic to thermophilic operation (**Table S1**). A total of more than 50,000 observations, including operational, physicochemical, and performance parameters, except volatile fatty acids (VFAs), were obtained from the records of individual plants. Each key variable had at least 1,087 observations, except VFAs (**Table 1**).

**Table 1.**
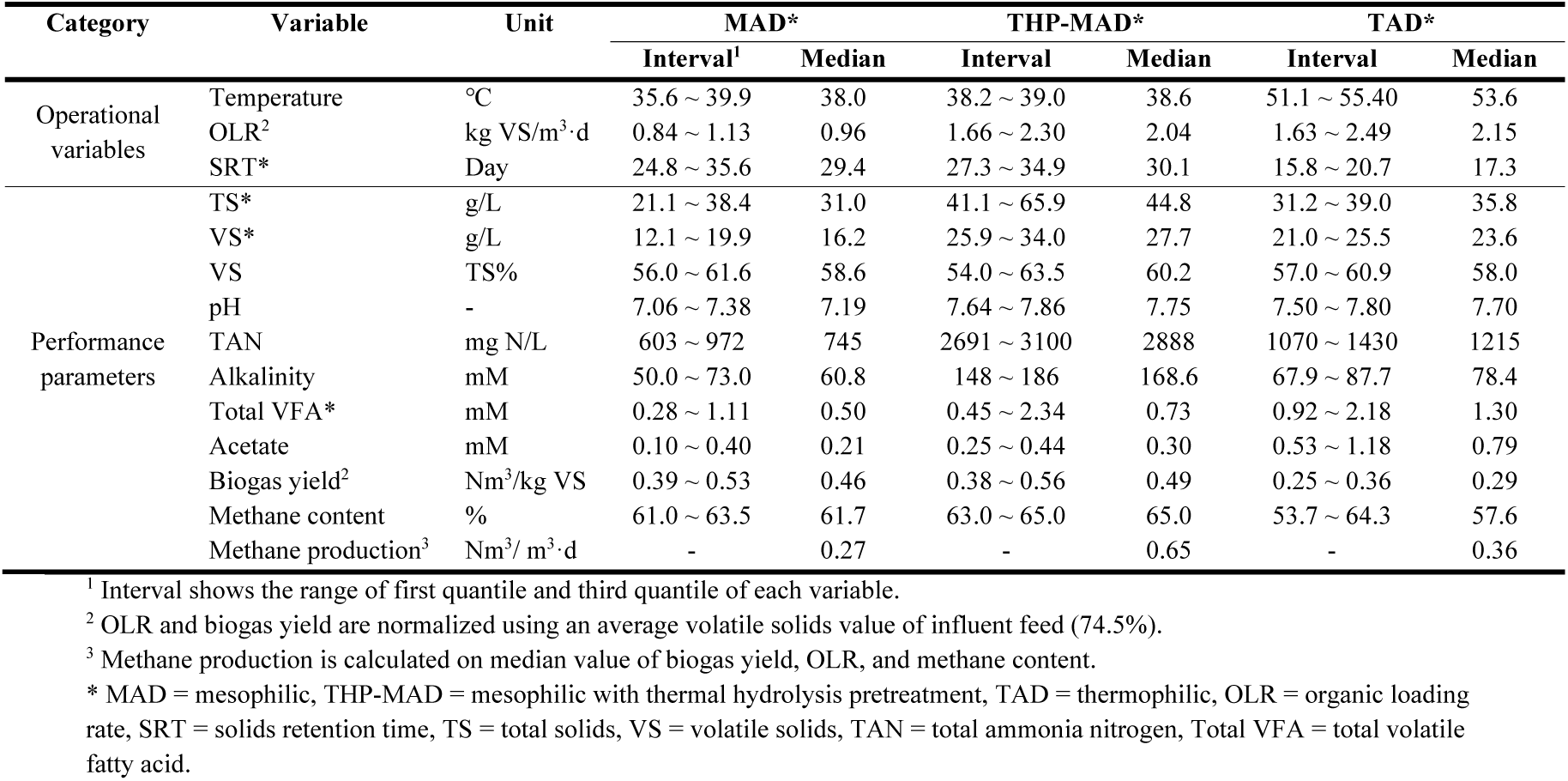
Operational variables and performance parameters: Intervals and median values for ADs at WWTPs in Denmark.

The digester sludge samples were obtained 2-4 times a year during the investigation period, and the VFA samples were collected 2-5 times from each studied digester during 2016. All samples were transported to the laboratory within 24 h and processed immediately upon arrival. After homogenization, the biomass samples were stored as 2 mL aliquots at -80°C before DNA extraction. Samples for VFA analysis were filtered with 0.22 µm filters (Frisenette, Knebel, Denmark) and stored at -20°C until analysis, which is described elsewhere [29].

### DNA extraction, 16S rRNA gene amplicon sequencing, and bioinformatics processing

The microbial communities of a total of 1,010 AD sludge samples (418 for archaea and 592 for bacteria) were analyzed using 16S rRNA gene amplicon sequencing. 50 µl AD sample were used for DNA extraction with the FastDNA® Spin Kit for soil (MP Biomedicals, Solon, OH, USA), following the optimized protocol for anaerobic digesters by Kirkegaard et al. [30]. The library preparation for 16S rRNA amplicon sequencing was performed as described in Kirkegaard et al. [10], targeting the V1-3 variable regions for bacteria and V3-5 variable regions for archaea. The bacterial primers used were 27F (AGAGTTTGATCCTGGCTCAG) [31] and 534R (ATTACCGCGGCTGCTGG)[32], which amplify a DNA fragment of ∼500 bp of the 16S rRNA gene (V1–3). The archaeal primers used were 340F (CCCTAHGGGGYGCASCA) [33] and 915R (GWGCYCCCCCGYCAATTC) [33], which amplify a DNA fragment of ∼ 560 bp of the 16S rRNA gene (V3–5). The amplicon libraries were paired-end sequenced (2×300 bp) on the Illumina MiSeq as described by Albertsen et al. [34].

The archaeal and bacterial read data were analyzed separately using USEARCH (v.11.0.667) [35]. For the V1-3 amplicons raw fastq files were filtered for phiX sequences using -filter_phix, trimmed to 250 bp using -fastx_truncate -trunclen 250, and quality filtered using -fastq_filter with -fastq_maxee 1.0. The sequences were dereplicated using -fastx_uniques with -sizeout -relabel Uniq. ASVs were generated using UNOISE3 [36], and ASV-tables were created by mapping the raw reads to the ASVs using -otutab with the -zotus and -strand both options. Taxonomy was assigned using the MiDAS 3 reference database [24,25] using sintax with the -strand both and -sintax_cutoff 0.8 [37]. The V3-5 amplicon data were analyzed in the same way except that only the reverse read was used and the primer binding site was removed during the trimming using –fastx_truncate – stripleft 18 –trunclen 250.

### Data processing and statistical analysis

Downstream statistical analyses and visualization were mostly performed in the R environment (v3.6.2) [38] using *ampvis2* (v2.5.8) [34] and *ggplot2* (v3.2.1) [39], unless indicated otherwise. Non-parametric *dunn*.*test* was used to identify significant differences between AD types. The correlations between all the variables were explored by Spearman correlation, where correlations greater than ±0.5 and false discovery rate (FDR) corrected *P* > 0.05 were visualized in Gephi (v0.9.2) [40], using Force Altas2 and manual tweaking to generate the network. For sequence data, samples were randomly subsampled to 10,000 sequences per sample, yielding a final dataset of 402 archaeal and 564 bacterial samples. For the growing bacteria datasets, after removing the non-growing ASVs from ASVtable, samples were also randomly subsampled to 10,000 sequences per sample for downstream analysis and comparison. Boxplot and heatmaps were made by the *amp_boxplot* and *amp_heatmap* function in *ampvis2*. Alpha diversity was calculated by *amp_alphadiv* function in *ampvis2*. The linear regression between alpha diversity (using Shannon’s index) and each operational and performance variable was used to pick the key variables most correlated. Weighted uniFrac distance, calculated by *beta_diversity*.*py* script in QIIME (v1.9.0) [41], was applied for all beta diversity comparisons. For ordination visualizations, the non-metric multidimensional scaling (NMDS) was performed by *amp_ordinate* in ampvis2 to show the dissimilarities of microbial profiles. Based on weighted uniFrac distance matrix, ANOSIM was applied to assess similarities for categorical variables using *compare_categories*.*py* in QIIME with 999 permutations. A PERMANOVA analysis using adonis in QIIME was used to describe the strength and significance for continuous variables. The significant difference of species between two groups of feed sludge was explored by Wilcoxon rank-sum test.

PLS regression was performed using R package *mdatools* v0.10.1 [21] to validate quantitative relationship between operational and performance parameters, and the microbial community, as well as to identify specific microbes which correlate to each variable the most. All bacterial species with median relative abundance >= 0.01% and archaeal ASVs with median relative abundance >= 0.05% were used to perform the PLS analysis. The model was trained using all samples and validated by segmented cross-validation (CV) with systematic splits (venetian blinds). Determination coefficient (R^2^) and root mean square error (RMSE) were used to assess performance of the model. The contribution of individual predictors was evaluated using regression coefficients and corresponding inferential analysis carried out by Jack-Knifing approach [42].

## Results and discussion

### Characterization of key parameters of AD

Key operational and performance parameters of the 46 anaerobic digesters during the six-year survey are summarized in **Table 1**. The digesters are classified into three types, based on the operational temperature and pretreatment of the feed sludge. MAD is the most common configuration (78% of all digesters) followed by TAD (15%) and THP-MAD (7%). The most common digester type is single-stage continuously stirred tank reactor (CSTR). The anaerobic digesters surveyed were running stably without major process complications for six years, therefore common ranges of operation and performance conditions are described for each digester type. As presented in **Table 1** and described below, values of several environmental parameters are very different from other AD systems treating manure, crops, food waste, and industrial waste [7,11,15,43–47], with generally lower or much lower values of total ammonia nitrogen (TAN) and level of VFAs.

The median temperature values of the three types of digesters were 38.0°C, 38.6°C, and 53.6°C, for MAD, THP-MAD, and TAD, respectively. Other operational variables (OLR and SRT) and the performance parameters (pH, TAN, alkalinity, TS, VS, biogas yield, and methane content) were found to be significantly different across all three types of AD (**Figure S2**). For more details on the description of each parameter, please see **Additional file 2**. In general, the same overall correlations between operational and performance parameters across digesters treating different types of substrates could also be observed specifically for digesters among WWTPs. Strong positive correlations (Spearman, r > 0.7, false discovery rate (FDR) *P* < 0.05) were observed between TAN and TS or VS, OLR and alkalinity, and methane production and SRT (**Figure S3**). Strong negative correlations between the OLR and methane production and biogas yield were also revealed (Spearman, r > 0.65, FDR *P* < 0.05), and between methane production and TAN (**Figure S3**), indicating that these operational variables are linked to the performance of the digesters. It is interesting that VFAs only related weakly to SRT and the ratio of VS to TS, which have previously been considered as important variables [48]. This may be due to a general low concentration range in the digesters and low loading.

Like many other full-scale plants, running at low OLR and long SRT of the digesters surveyed, which is referred as “suboptimal” operational conditions, may lead up to a 30% profitability loss [46,49]. Increasing the OLR seems promising, but there may be a number of operational problems which need to be considered, such as foaming and acidosis, due to the imbalance between operational and microbial processes. Thus, a better understanding of microbial communities and their function may help to control or manipulate the processes that decrease the potential risks of operational failures.

### Bacterial and archaeal microbiomes

We obtained 33,047 bacterial and 878 archaeal unique ASVs, which were classified and assigned using sintax and the MiDAS 3 database. Thus, a total of 42 phyla, 1,600 genera, and 3,584 species were detected in the bacterial microbiome, where 1,117 (70%) genera and 3,336 (93%) species were novel and could only be assigned genus and species name based on the MiDAS 3 *denovo* placeholder taxonomy. For archaea it was not possible to analyze the methanogenic archaea at the species level because the phylogeny of most of these species cannot be resolved using the 16S rRNA gene, even with full-length sequences [24]. As a result, only 26 species were classified and most of the archaeal population are shown at ASV level.

AD at WWTPs are complex systems, as they receive a substantial amount of microorganisms via feed streams (primary sludge, PS, or surplus activated sludge, AS). Many of these microorganisms are not growing in the digester, presumably inactive or dying off [10,13,50]. The growing and non-growing microorganisms were identified according to the ratio of read abundances in digester and feed as described by Kirkegaard et al. [10]. The bimodal distribution of ratios was split at a ratio of around 10 (**Figure S4**), showing two clearly separated groups of ASVs. The group with a ratio >10 represents ASVs enriched in digesters compared to the feed sludge, here designated as “growing microorganisms”. The group with a ratio <10 represents ASVs with unchanged or lower relative read abundance in digesters, compared to the feed sludge, here designated as “non-growing microorganisms”. Thus, combined with median relative abundance across samples in each type of AD, the total ASVs (>0.01% median relative abundance) were divided into four groups, growing/non-growing ASVs with high abundance (>0.1%) and growing/non-growing ASVs with low abundance (<0.1%). It was observed that the growing highly abundant ASVs only accounted for 7.6%, 23.2%, and 9.4% of the total ASV counts in MAD, THP-MAD, and TAD, respectively (**Fig. 1**). However, the proportion of relative abundances of these growing highly abundant ASVs were large, at 38.8%, 85.3%, and 50.9% in MAD, THP-MAD, and TAD, respectively. This suggests that the performance and functionality of AD might be driven by only a small number of the microbial phylotypes detected by amplicon sequencing.

**Fig. 1.**
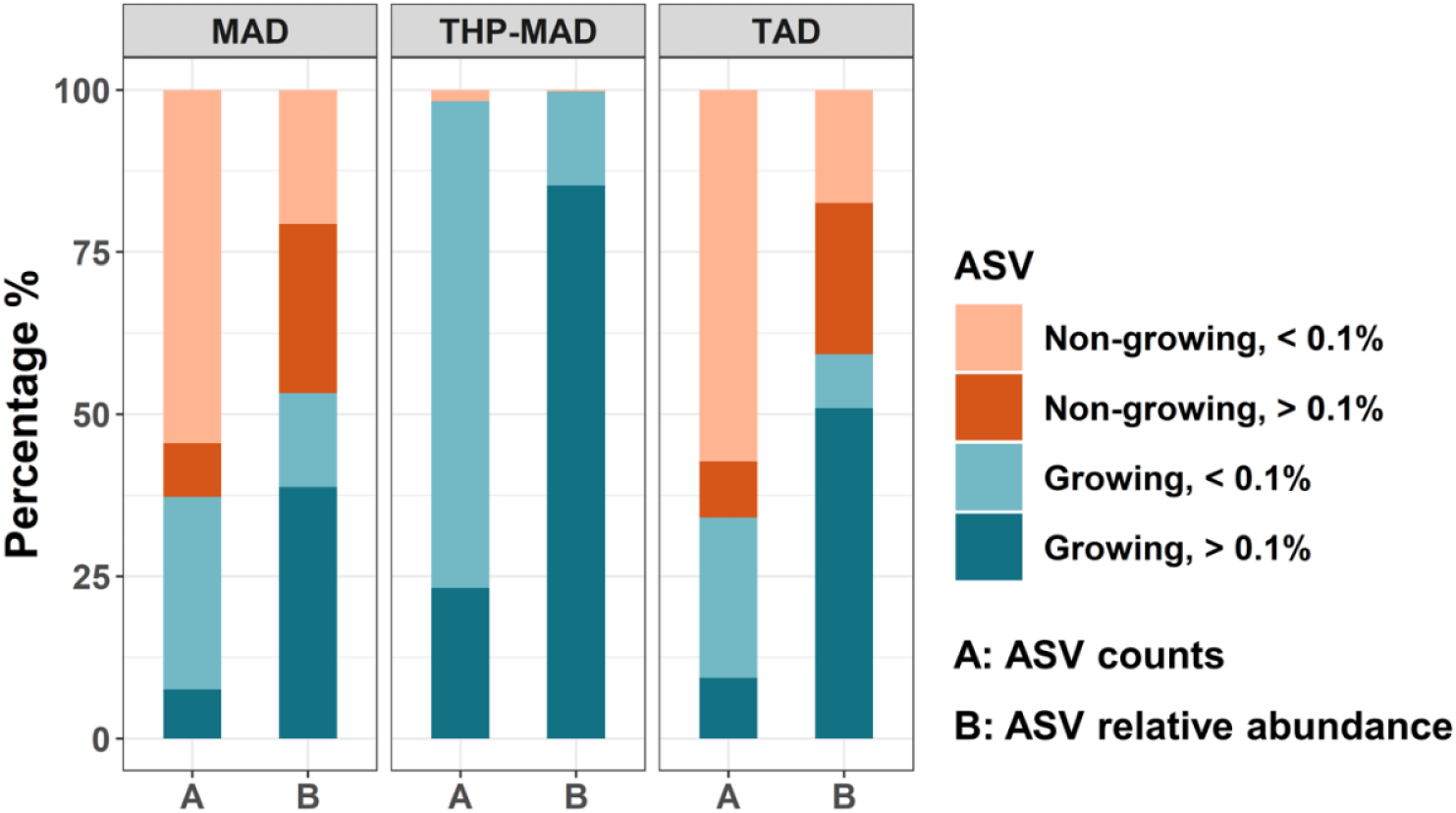
Composition of growing and non-growing ASVs in Danish ADs at WWTPs. The total ASVs (median relative abundance > 0.01%) divided into four groups based on growth ratio (growing/non-growing) and relative abundance (high/low abundant, 0.1% indicates the cutoff value). A and B show the composition of ASV counts and ASV relative abundance for these four groups in MAD, THP-MAD, and TAD, respectively. MAD = Mesophilic AD, THP = Mesophilic with thermal hydrolysis pretreatment, TAD = Thermophilic AD.

The five most abundant bacterial phyla were Firmicutes, Proteobacteria, Chloroflexi, Actinobacteria, and Bacteroidetes, accounting for 75.7% (median value) of all amplicon sequences across all samples, and these phyla are typical for digesters at WWTPs [7,15,43,51–54]. However, the three types of AD showed variations in the dominant bacterial taxa, especially at genus level (**Figure S5**). Among the 25 most abundant species, 11 species in MAD and 9 species in TAD belonged to the group of non-growing microorganisms (**Fig. 2A** and **2B)**. These included species in genera belonging to the polyphosphate-accumulating organisms (PAO) Tetrasphaera, the putative PAO *Dechloromonas* [55], the filamentous genus *Ca*. Microthrix [56], and the genera *Romboutsia*, and *Trichococcus*. These all belong to the top-most abundant reported genera in activated sludge in Danish WWTPs [25], thereby indicating carry-over to the digesters with the feed sludge. Since the top100 species in MAD and TAD, respectively, are very similar across the digesters, these lists can be used as a representative reference of abundant growing and non-growing organisms in digesters at WWTPs across the world **(Figure S6 and S7)**. These results demonstrate that surprisingly many, 44% and 54% of the species were non-growing in MAD and TAD, respectively. The top 25 species in THP-MAD all belonged to growing microorganisms in good agreement with the presence of THP pretreatment, which causes a decay of essentially all organisms coming with the feed sludge (**Figure S8A**).

**Fig. 2.**
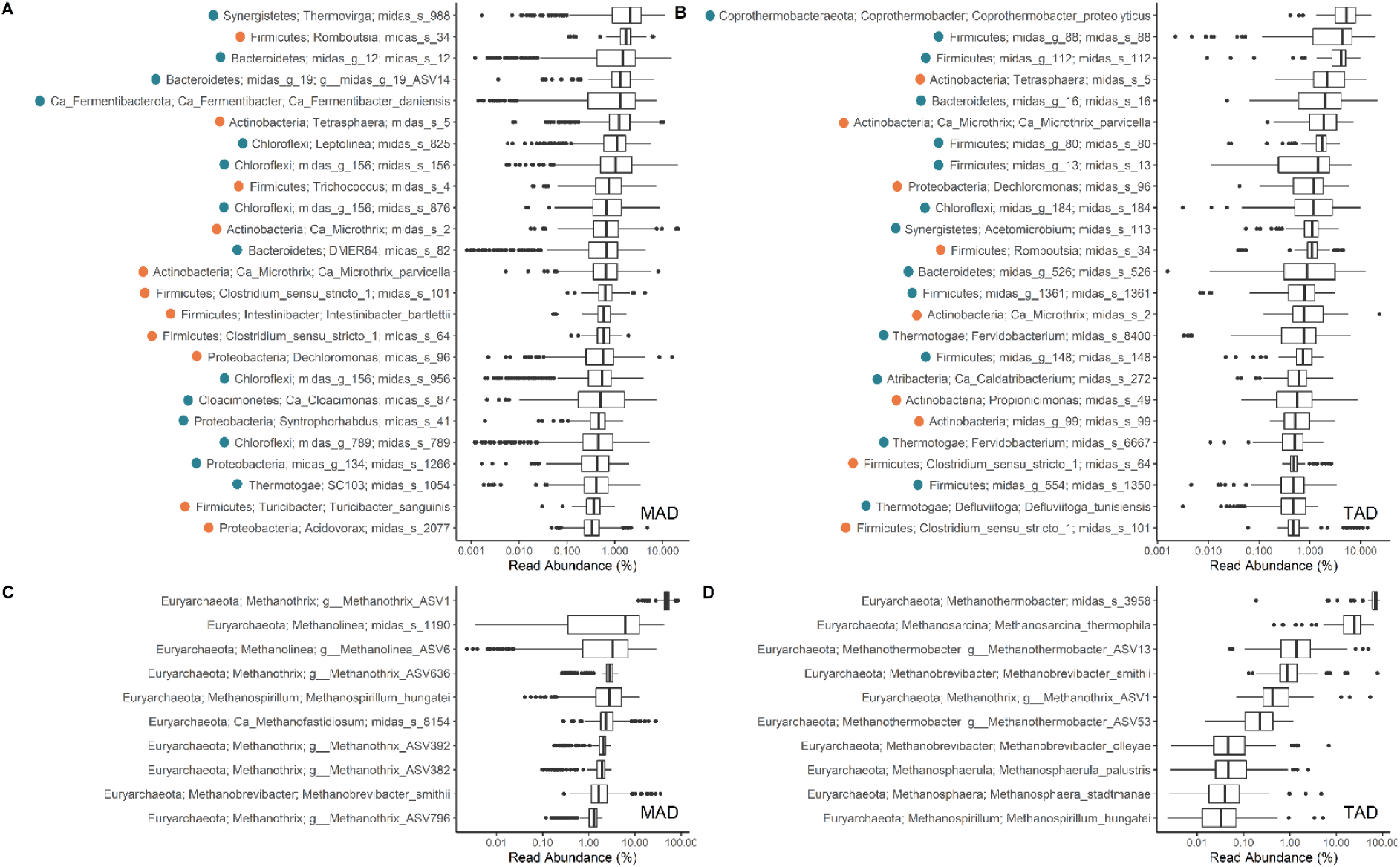
Boxplots of the most abundant species/ASVs in Danish ADs at WWTPs. (A) The 25 most abundant bacterial species/ASVs in MAD, (B) The 25 most abundant bacteria species/ASVs in TAD, (C) The 10 most abundant archaeal species/ASVs in MAD, (D) The 10 most abundant archaeal species/ASVs in TAD. The dots at the left in A and B indicate whether the species/ASVs are growing (ratio >10, blue), non-growing or dying off (ratio <10, orange). MAD = Mesophilic AD, TAD = Thermophilic AD. Ratio refers to the digester to influent relative read abundance ratio (please see **Additional file 3**). Sequences not possessing species-levels classification are shown here as individual ASVs.

The growing microorganisms are considered to be responsible for the most important ecological functions within AD. Among the dominant growing bacterial species there were many known fermenters, such as species belonging to the genera *Thermovirga, Ca*. Fermentibacter, and *Leptolinea* in MAD, and *Coprothermobacter* and *Acetomicrobium* in TAD. There were also syntrophic bacteria, such as members of *Ca*. Cloacimonas [57] and *Syntrophorhabdus* in MAD. However, a large fraction of the most abundant growing species were novel taxa without any known function. They were identified by MiDAS 3 species-level taxonomy and given robust placeholder names until characterized in more detail, enabling across-study comparisons of AD microbiome at high taxonomic resolution [24]. Due to the high relative abundance, some genera are of special interest: midas_g_12 (family Prolixibacteraceae), midas_g_19 (family Bacteroidetes vadinHA17), midas_g_156 (family Anaerolineaceae), and midas_g_789 (family Anaerolineaceae) in MAD; midas_g_88 (family Syntrophomonadaceae), midas_g_112 (order MBA03), and midas_g_16 (family Lentimicrobiaceae) in TAD, and midas_g_13 (order D8A-2) in THP-MAD. Some of these genera encompass very abundant species, especially in the family Anaerolineaceae (up to 8% median abundance): midas_s_156, midas_s_876, midas_s_956, midas_s_1462, midas_s_467, midas_s_1625. These abundant and novel taxa should be investigated in future studies, as their physiology and ecological role in AD are completely unknown while likely important.

Moreover, compared with MiDAS 2 taxonomy (which was the curated version of Silva taxonomy) [58], MiDAS 3 provides a much higher resolution to classify sequences and introduces species-level names for the first time for the abundant microorganisms in AD ecosystem. For example, genus T78 (family Anaerolineaceae) in MiDAS 2 encompassed sequences that are split into midas_g_156 and midas_g_467 in MiDAS 3, both being the abundant genera mentioned above. These genera are both diverse, each having three abundant species present in MAD (**Figure S9**). Also, *Ca*. Cloacimonas and *Pelotomaculum* and the newly discovered syntrophic genus midas_g_995 [29] had high species diversity as well, with several abundant species with random distribution (**Figure S9**).

Euryarchaeota was the dominant archaea in the digesters (99.9%, median value). The acetoclastic genus *Methanothrix* (previously named *Methanosaeta*) dominated in MAD (71.8%) and THP-MAD (93.8%), whereas the genera *Methanothermobacter* (70.7%) and *Methanosarcina* (24.8%) dominated in TAD (**Figure S10B**). *Methanosarcina* was in very low abundance in MAD (0.1%) and THP-MAD (0.01%), in contrast to other mesophilic full-scale studies of manure-based AD where it was dominant [7,53,59]. However, the low concentrations of VFAs (<1 mM) in the mesophilic AD may explain why *Methanothrix* dominated [60]. The hydrogenotrophic methanogenic *Methanoculleus* was the predominant genus (4.2%) in THP-MAD, which is in line with other lab-scale and pilot-scale THP digesters studies [61,62].

Acetoclastic and hydrogenotrophic methanogenic species/ASVs were quite abundant in all the types of AD (**Fig. 2C, 2D, and Figure S8B**). g_*Methanothrix*_ASV1 was dominant in both MAD and THP-MAD, followed by midas_s_1190 (genus *Methanolinea*) and g_*Methanolinea*_ASV6 in MAD and midas_s_880 (genus *Methanoculleus*) in THP-MAD. *Methanothermobacter tenebrarum* and *Methanosarcina thermophila* species were the second most abundant species in TAD (**Fig. 2**).

### Microbial diversity in different AD types

The use of common measures for richness and diversity in the digesters surveyed has only limited value, as abundant immigrating bacteria, likely inactive or dying off without any functional role in the systems, will influence the diversity measures and produce misleading results. This is illustrated by comparing the diversity measures calculated for all bacteria and for the growing bacteria only (**Fig. 3A**). When the non-growing fraction was removed, the median values of observed ASVs decreased from 1935 to 928, 1486 to 534, and the median values of Shannon index from 6.22 to 5.11 and 5.59 to 4.33 in MAD and TAD, respectively (**Fig. 3A**). THP-MAD only had a minor change in observed ASVs (from 832 to 741) and the median Shannon index (4.47 and 4.63, respectively), reflecting, as expected, that these communities were not strongly influenced by immigration. The adjusted diversity measures showed the same order of magnitude decrease for archaea (**Figure S11**) with THP-MAD between MAD and TAD. The measures also showed that higher temperature harbored fewer number of microbes in accordance to other full-scale surveys [15,51], but that the exact values are strongly dependent on the inclusion of the immigrating microbes. Higher alpha diversity measures for bacterial communities compared with archaea is in agreement with other full-scale WWTPs studies [15,43,51]. The diversity in thermophilic AD has been shown to be lower than in mesophilic digesters [63–65], which is also supported by our data.

**Fig. 3.**
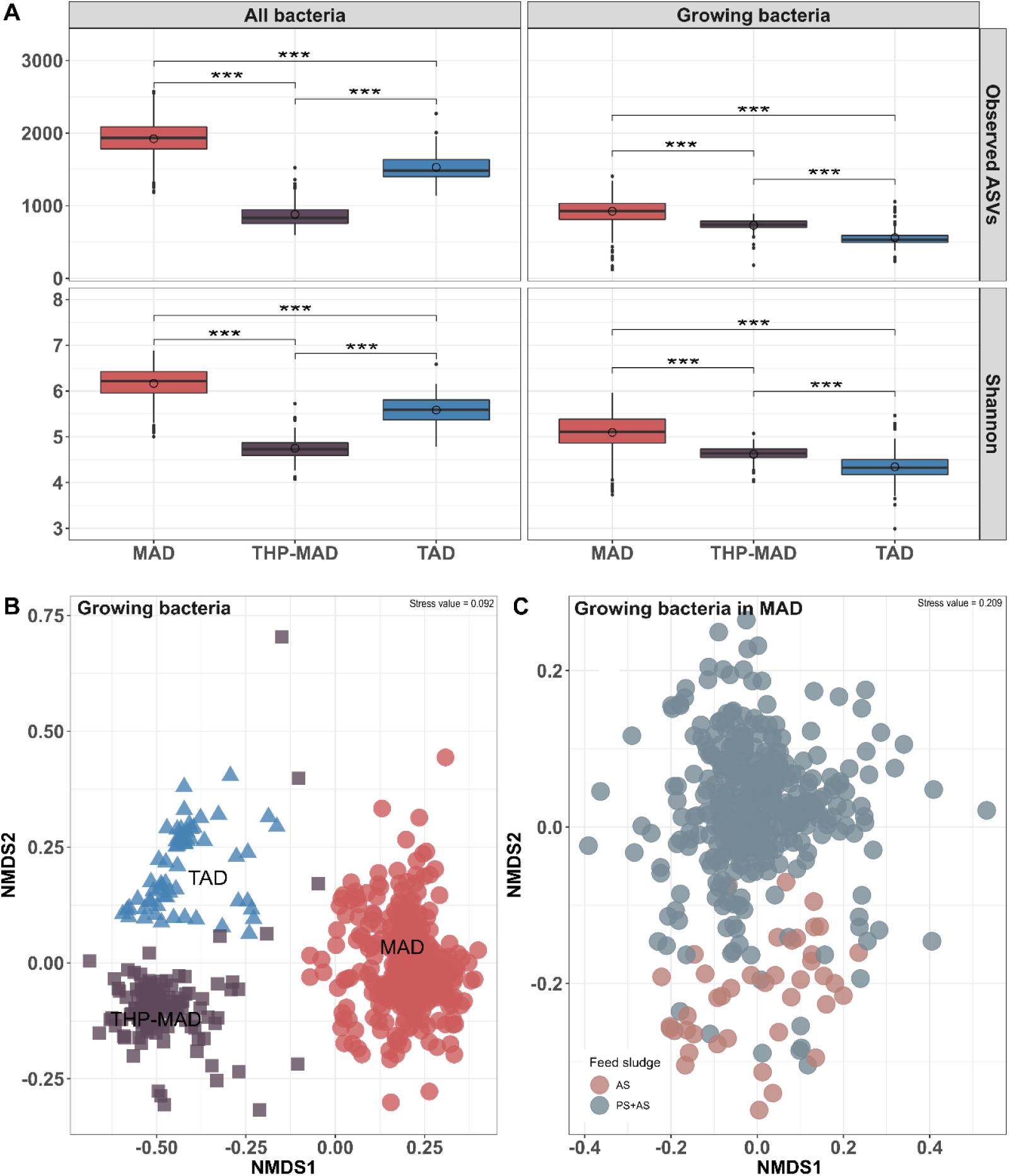
Alpha and beta diversity plots of bacterial and archaeal communities of three types of AD. (A) Boxplots of observed ASVs and Shannon index of entire and growing bacterial community, significant differences are indicated (Wilcoxon rank-sum test; ***, p < 0.001). (B) Non-metric multidimensional scaling (NMDS) plots of growing bacterial community structure based on weighted UniFrac matrix, (C) NMDS plots of growing bacterial community structure of MAD based on weighted UniFrac matrix. MAD = Mesophilic AD, THP-MAD = Mesophilic with thermal hydrolysis pretreatment process, TAD = Mesophilic AD, AS = Activated sludge, PS = Primary sludge.

The total bacterial (including growing and non-growing fraction) and archaeal microbiome seemed relatively stable in each digester across all 22 WWTPs during the six-year survey as indicated by tight clustering as visualized by non-metric multidimensional scaling (NMDS) (**Figure S12**). This is also reported from other time-series studies of full-scale digesters mainly treating manure, agricultural waste, and municipal solid waste [7,43], suggesting that overall stable microbiomes are common in full-scale digesters during steady-state operation. However, as a major part of the microorganisms are immigrants, they may strongly affect the betadiversity measures. Therefore, it is important to compare the diversity of both the total and the growing fraction of the population. The dissimilarity among plants seemed the same considering the community structure of the total and growing bacteria in MAD (ANOSIM; Total bacteria: R = 0.65, *P* = 0.001; Growing bacteria: R = 0.63, *P* = 0.001) and THP-MAD (ANOSIM; Total bacteria: R = 0.45, *P* = 0.001; Growing bacteria: R = 0.49, *P* = 0.001). However, the growing bacterial community in TAD became more similar across plants compared to the total bacterial population (ANOSIM; Total bacteria: R = 0.64, *P* = 0.001; Growing bacteria: R = 0.40, *P* = 0.001). This shows that the inclusion of non-growing bacteria in microbiome analyses of TAD may lead to misleading results and erroneous conclusions.

Analysis of beta diversity of the communities present in different AD types revealed three distinct clusters corresponding to MAD, THP-MAD, and TAD (**Fig. 3B and Figure S13A**). Clear separation dictated by AD type was evident for all bacteria (ANOSIM: R = 0.95, *P* = 0.001), the growing bacteria (ANOSIM: R = 0.97, *P* = 0.001), as well as the archaeal microbiome (ANOSIM: R = 0.83, *P* = 0.001), reflecting the huge effects of operational conditions on the resulting variation in the microbiomes. Permutational multivariate analyses of variance showed that TAN contributed to shaping the structure of the total bacterial microbiome (adonis: R^2^ = 32%, *P* = 0.001) (**Figure S13B**), which has also been observed in full-scale digesters treating different kinds of substrates [7]. In contrast to the bacterial microbiome, the overall structure of the archaeal microbiome was separated mainly by temperature (adonis: R^2^ = 66%, *P* = 0.002), with a separate cluster for THP-MAD alongside MAD (**Figure S13C**). pH was the second factor influencing the archaeal microbiome (adonis: R^2^ = 27%, *P* = 0.002), which may explain the separated cluster of THP-MAD from MAD (**Figure S13D**).

### Main drivers of MAD microbiome

Since most digesters surveyed were MAD, we further applied the correlation analysis between key parameters and microbial diversity and structure to determine the main drivers, with special focus on the growing bacterial microbiome, as non-growing microorganisms may mask the influence of key drivers on the active microbiome in correlation analyses. In general, bigger difference was observed on linear regression of key parameters against alpha diversity between the total and growing bacterial microbiomes, compared with permutational multivariate analysis of betadiversity (**Table S2** and **Table S3**).

It is well-known that temperature is a very important factor for shaping the microbial diversity and community structure in full-scale digesters [7,15,43], but it is less clear to what extent it is for mesophilic AD at WWTPs. In our study, the temperature range in MAD was small (35.6-39.9) and was only considered to be most important to the total bacterial alpha diversity in MAD (25%, linear regression, FDR *P* < 0.001, **Table S2**), but not the alpha diversity of growing bacteria (16%, FDR *P* < 0.001). This indicates that temperature may not be the most important factor in MAD. Instead, the correlation coefficient of OLR improved significantly by subsetting the growing bacterial alpha diversity (31%, *P* < 0.001) compared to the total bacterial alpha diversity (9%, *P* > 0.05). OLR also shaped the microbiome structure (beta diversity) of growing bacteria (adonis: R^2^ = 21%, *P* = 0.001, **Table S3**). Although OLR is widely accepted as a deterministic factor for any type of AD microbiome [66–69], our study strengthened it when OLR was only correlated with growing bacteria. Moreover, the biogas yield exhibited strong correlation with the growing bacterial microbiome both on alpha diversity (46%, *P* < 0.001) and beta diversity (adonis: R^2^ = 31%, *P* = 0.001), as well as archaeal beta diversity (adonis: R^2^ = 23%, *P* = 0.008), supporting the observation that AD performance depends on the activity of the microbiome [70]. Similarly, TAN was observed to be more correlated to growing bacterial alpha diversity compared with the total bacterial population (**Table S2**). Regarding the archaeal microbiome, no strong correlation was found between parameters and alpha diversity in MAD. However, apart from biogas yield, acetate concentration (adonis: R^2^ = 18%, *P* = 0.04) was also found to have significant correlation with the archaeal microbiome structure.

Samples from MAD digesters treating only surplus AS and without PS (Fornaes, Mariagerfjord, and Soeholt) formed a small separate cluster, compared to MAD treating a mixture of both types of substrates for all bacteria (ANOSIM: R = 0.44, *P* = 0.001, **Figure S14A**) and for the growing bacteria (ANOSIM: R = 0.57, *P* = 0.001, **Fig. 3C**). The higher dissimilarity of growing bacteria in MAD supports the observation that substrate characteristics (e.g., biodegradability, composition, concentration) from PS shape the growing bacterial community structure [7,15]. This is different from the growing bacteria in the three types of AD, which is mainly driven by operational parameters (**Fig. 3B**). While for all bacterial communities, the relative abundance of 18 of the 25 most abundant species showed a significant difference between the two clusters depending on the feed sludge (Wilcoxon rank-sum test, *P* < 0.05, **Figure S14B**). Non-growing species in digesters but abundant in AS (i.e., species belonging to genus *Tetrasphaera, Ca*. Microthrix, or *Dechloromonas*) were found in higher relative abundance in MAD treating only AS compared to MAD treating AS and PS. These results underline that besides substrate characteristics, the immigrating bacterial load has a strong impact on the total bacterial microbiome in AD. Additionally, we also observed the influence of feed sludge on archaeal community (**Figure S15A**), with significant difference between the digesters with different feed sludge (ANOSIM: R = 0.23, *P* = 0.001). It is interesting to see that the species belonging to genus *Methanolinea* were rare in the digesters only fed with AS (**Figure S15B**).

### Relationship between MAD microbiome and its driving factors predicted by PLS regression model

We applied PLS regression to predict key operational variables and performance parameters and their relationship to the microbiome in MAD. Separate PLS models were built on the bacterial microbiome at species level and archaeal ASVs with each factor (for bacteria: temperature, OLR, TAN, and biogas yield; for archaea: temperature, TAN, acetate, and biogas yield). Very good prediction accuracy was observed on the bacterial microbiome, where the CV R^2^ of all four PLS models exceeded 0.85 (**Fig. 4**). However, none of the models based on archaeal ASVs had the R^2^ over 0.80, except for biogas yield (**Fig. 4**). PLS regression models were also carried out on pH and SRT, since they are important AD parameters (**Figure S16 and S17**). The growing bacterial species and archaeal ASVs, which most significantly contributed to each PLS regression model (the contribution was estimated using inferential statistics for corresponding regression coefficients, *P* < 0.05), for both positive and negative contributions, are shown in **Fig. 5**.

**Fig. 4.**
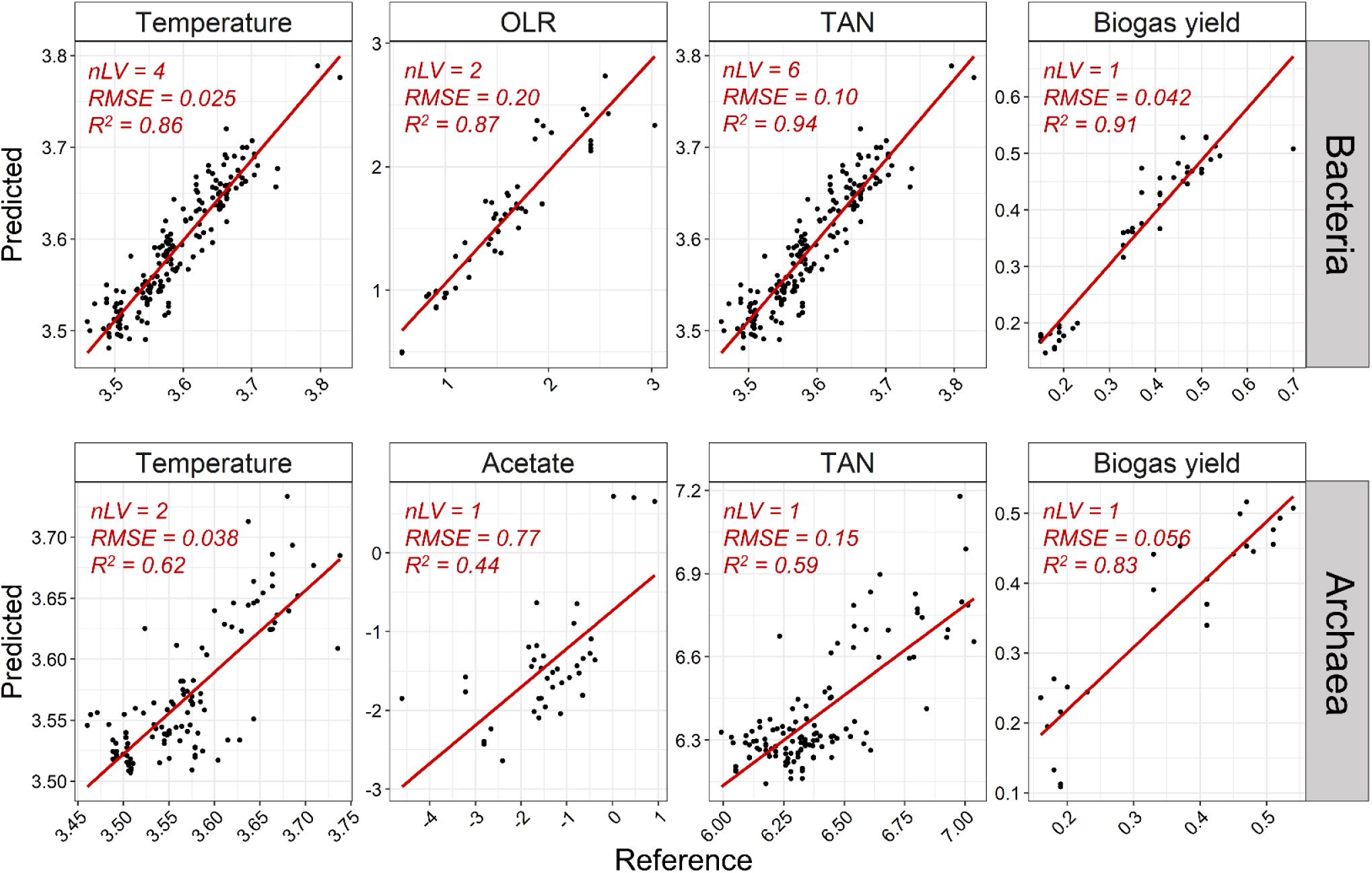
Prediction plots of main drivers based on bacterial and archaeal microbiome in MAD by partial least squares regression. MAD = mesophilic AD, OLR = organic loading rate, TAN = total ammonium nitrogen. nLV = number of selected components, RMSE = root mean squared error.

**Fig. 5.**
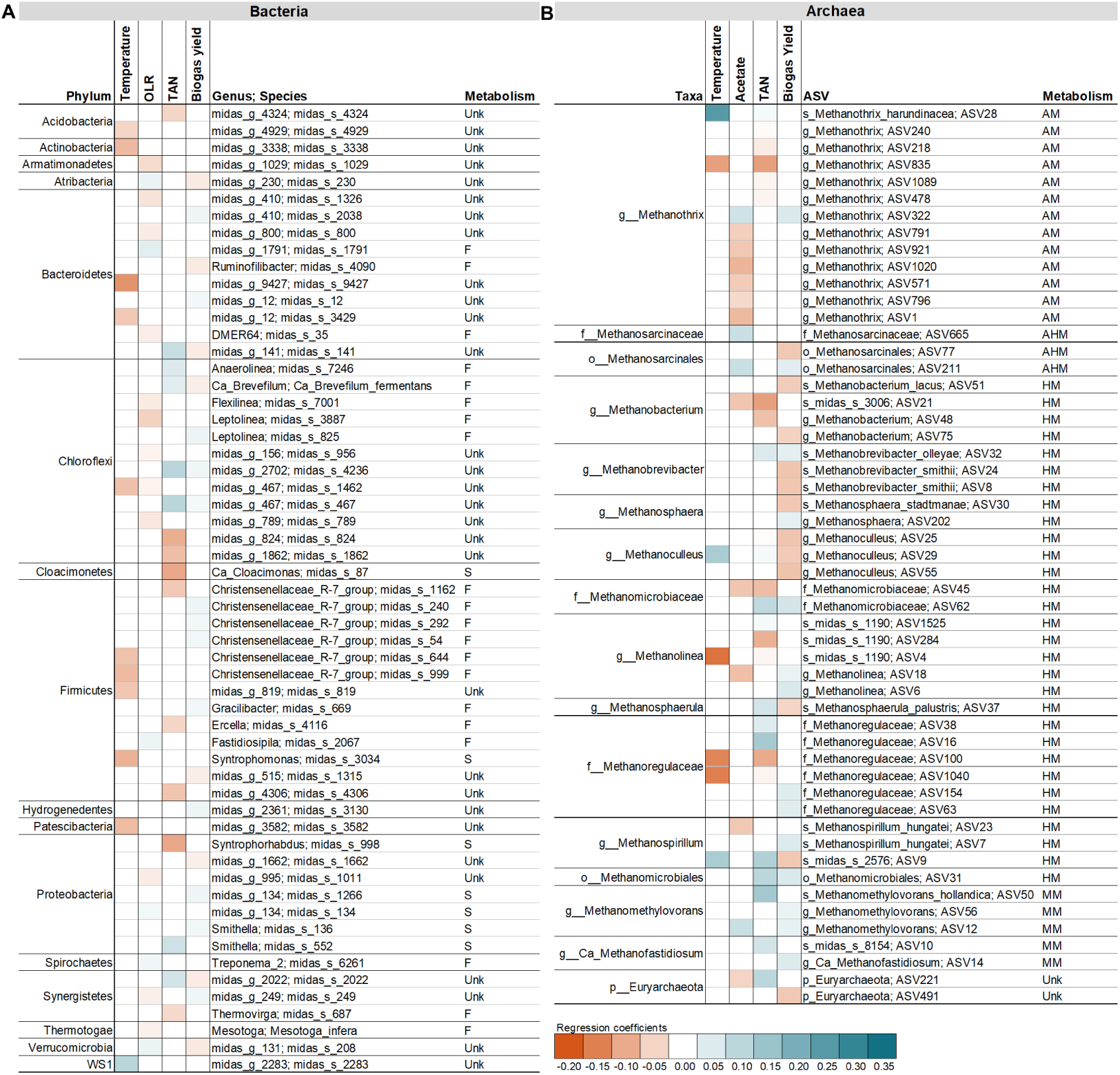
Partial least squares estimation for main driver for important growing bacterial species (A) and archaeal ASVs (B) in MAD. *P* < 0.05, positive correlation in blue, negative correlation in orange. F = Fermenters, S = Syntrophic bacteria, AM = Acetoclastic methanogens, AHM = Acetoclastic or Hydrogenotrophic methanogens, HM = Hydrogenotrophic methanogens, MM = Methylotrophic methanogens, Unk = Unknown.

Most growing bacterial species exhibiting significant correlations were represented by novel taxa (**Fig. 5A)**. Species belonging to the same genus were correlated to different operational or performance parameters, such as was the case with two species in family Anaerolineaceae (midas_s_467 and midas_s_1462 belonging to genus midas_g_467). The positive correlation of midas_s_467 with TAN and biogas yield, and the negative correlation of midas_s_1462 with temperature and OLR, could explain the abundance variability and trend across MAD (**Figure S9**). Similar observations were found for three species belonging to genus midas_g_12 (family Prolixibacteraceae, phylum Bacteroidetes). The ecological function of these novel species is unknown so the PLS correlation results may provide hypotheses which could aid the design of experiments to reveal the role of novel taxa in AD [71]. Among the known species, *Ca*. Brevefilum fementans showed a positive correlation with TAN, which can indicate a preference or tolerance to slightly higher TAN conditions. This hypothesis is supported by their genome blueprint indicating that *Ca*. B. fementans can ferment proteinaceous substrates to VFAs with ammonium as a by-product of protein degradation [72]. Additionally, species belonging to the known syntrophic genera *Ca*. Cloacimonas, *Smithella, Syntrophomonas*, and *Syntrophorhabdus* were found mostly negatively correlated with TAN and temperature, thereby confirming the high sensitivity of this group to environmental conditions [73–76].

Non-growing bacterial species (primarily immigrating with the feed sludge) showed negative correlations to some key parameters (**Figure S18**), especially for biogas yield and SRT, suggesting that they are not directly involved in the conversion of feed stocks to biogas and are probably degraded or washed out of the digesters. This is exemplified by *Tetrasphaera* midas_s_5, the most abundant non-growing species in MAD, and it was negatively correlated with SRT together with *Tetrasphaera elongata* and midas_s_299. *Tetrasphaera* is very abundant in Danish wastewater treatment plants [55] and is introduced into the digester with the waste activated sludge. The negative correlation with the SRT indicates that *Tetrasphaera* is dying off in the digesters despite the potential for surviving or growing under anaerobic conditions as fermenters, and polyphosphate accumulators [77].

Correlation results for archaeal ASVs are shown in **Fig. 5B**. Generally, ASVs belonging to the same genus show the same trends. For example, ASVs classified to family Methanosarcinaceae and order Methanosarcinales known as acetoclastic and hydogenotrophic methanogens, positively correlated with acetate. This is consistent with other findings [78] where *Methanosarcina* was most abundant in digesters with higher acetate concentration. In contrast, most *Methanothrix* ASVs correlated negatively with acetate, supporting the dominance of *Methanothrix* at low acetate concentrations [79]. Many *Methanothrix* ASVs also showed negative correlation with TAN, which is in agreement with studies that show that *Methanothrix* is the methanogen most sensitive to ammonia inhibition [80]. It is interesting that even small TAN variations as seen here for Danish digesters (603-972 mg N/L, MAD – **Table 1**) can affect individual *Methanothrix* ASVs in different ways.

For hydrogenotrophic methanogens, different ASVs from the same genus or species showed diverse correlations with AD parameters. The second most abundant archaeal species in MAD belonging to genus *Methanolinea* (midas_s_1190) included three ASVs, which correlated with TAN differently suggesting that species microdiversity can influence process performance. The negative correlation of many hydrogenotrophic methanogens with biogas yield, such as *Methanoculleus*, could be due to their ability to survive/compete at sub-optimal AD conditions (e.g., increased TAN) [80] which could translate into lower biogas yields.

Overall, the PLS regression models presented enable elucidation of the relationships between important AD parameters and the main drivers shaping AD microbiome at very high resolution. The results for known taxa agree with present knowledge, thus verifying the robustness of the PLS application in microbiome study. Importantly, a combination of the PLS regression with species-level microbial data provides the first insight into potential functional importance of several novel microorganisms, where little or no description of their ecology and physiology is available. Based on our observations, hydrolytic-fermentative bacteria, and acetogenic syntrophs along with archaeal methanogens, all have significant and quantitative relationships with important parameters at MAD. This shows great promise for the improved models, and optimized functional performance of AD.

## Conclusion

A six-year survey of 46 anaerobic digesters located at 22 Danish WWTPs provided a comprehensive overview of typical operational and performance parameters, detailed identification of the AD microbiome at species level, and elucidated relationships between specific taxa and key parameters in AD. The anaerobic digesters surveyed were running stably but operated at low intensity, a common feature across digesters in WWTP. Non-growing species migrating from the feed sludge were abundant in mesophilic and thermophilic AD, but did not seem to contribute to the functionality of AD. In contrast, many growing species were novel and identified using MiDAS 3 taxonomy, and their physiological and ecological roles in AD remains to be described. The microbiome of the three types of AD surveyed (mesophilic, thermophilic, and thermal hydrolysis pre-treatment-mesophilic) showed high stability within plants, forming separate clusters for all bacteria, growing bacteria, and archaea depending on the operational parameters. The variations of growing bacteria within mesophilic digesters were related to organic loading rate, ammonium concentration, feed sludge characteristics, and biogas yield. Multiple correlations between growing bacteria and archaea at species level and key parameters were found, forming a basis for future studies of the ecology and function of novel taxa.

## Supporting information

Additonal file 1: Supplementary tables and figures

Additonal file 2: Characterization of key parameters of ADs

Additional file 3: Digester to influent relative read abundance ratios for each ASV

## Supplementary information

**Additional file 1**: **Table S1**. Overview of WWTP digester capacities, type, and industrial load. **Table S2**. Linear regression of key variables individually against alpha diversity using the Shannon diversity index at MAD. **Table S3**. Permutational multivariate analysis (using continuous variables only) of variance of beta diversity using weighted UniFrac matrix at MAD. **Figure S1**. Comparison of classification between full-length exact sequence variants (FL-ESVs) database and the SILVA_132_SSURef_Nr99 database on the top 50 ASVs in the digester sludge samples from Danish wastewater treatment plants. **Figure S2**. Box plots of operational and performance parameters of three types of AD. **Figure S3**. Spearman correlations on operational and performance parameters in AD. **Figure S4**. Distribution of digester:feed relative read abundance ratios for each ASV. **Figure S5**. (A) Relative abundance of the 20 most abundant bacterial phyla in AD. (B) Relative abundance of the 50 most abundant bacterial genera in AD (n = 564). **Figure S6**. Boxplots of the top 100 species/ASVs in MAD. **Figure S7**. Boxplots of the top 100 species ASVs in TAD. **Figure S8**. Boxplots of the most abundant species/ASVs in THP-MAD. **Figure S9**. Heatmap of the most abundant species/ASVs belonging to the genus T78 in MiDAS 2 (split into the genera midas_g_156 and midas_g_467, all family Anaerolineaceae), genus *Ca*. Cloacimonas, genus *Pelotomaculum*, midas_g_995, and genus *Methanothrix* in Danish digesters at WWTPs. **Figure S10**. Relative abundance of the 25 most abundant archaeal genera in AD (n = 402). **Figure S11**. Boxplots of alpha diversity measures of archaeal community of three types of AD. **Figure S12**. Non-metric multidimensional scaling (NMDS) plots of bacterial and archaeal community structure based on weighted Unifrac matrix colored by WWTPs. **Figure S13**. Non-metric multidimensional scaling (NMDS) plots of all aacteria and archaea community structures based on weighted Unifrac matrix. **Figure S14**. (A) Non-metric multidimensional scaling (NMDS) plots of entire bacterial community structure based on weighted UniFrac matrix in MAD. (B) Heatmap of 25 most abundant bacterial species in MAD digesters depending on the composition of feed sludge (AS and PS). **Figure S15**. (A) Non-metric multidimensional scaling (NMDS) plots of archaeal community structure based on weighted UniFrac matrix in MAD. (B) Heatmap of 15 most abundant archaeal species in MAD digesters depending on the composition of feed sludge (AS and PS). **Figure S16**. Prediction plots of other important parameters based on bacterial and archaeal microbiome in MAD by partial least square regression. **Figure S17**. Complete list for partial least squares estimation of key variables with growing bacterial species (A) and archaeal ASVs (B) in MAD. **Figure S18**. Partial least squares estimation of key variables with non-growing bacterial species in MAD.

**Additional file 2: Characterization of key parameters of anaerobic digestions**

**Additional file 3: Digester to influent relative read abundance ratios for each ASV**

## Acknowledgements

We thank the operators at different WWTPs in Denmark for providing digester sludge samples and plant records. The Chinese Scientific Research Council is acknowledged for providing financial support to C. Jiang.

## Authors’ contributions

PHN and CJ conceived and designed the work. CJ, MP, and PHN wrote the manuscript. CJ, MP, SK, and EY performed bioinformatic analysis, statistical analysis, and data visualization. KSA and MSD provided bioinformatics support. MHA, MN, RHK, and LH performed data collection, sampling, and lab procedures. JH and AAH contributed to sample and plant record collection. CJ, MP, PHN, SK, MN, and MSD contributed to data interpretation. All co-authors read and approved the final manuscript.

## Availability of data and materials

The raw amplicon sequences generated and analyzed during the current study were uploaded to the NCBI Sequence Read Archive (SRA) data depository, with the project number PRJNAXXXXX. Additional data can be shared upon request.

## Ethics approval and consent to participate

Not applicable

## Consent for publication

Not applicable

