## Supplementary material for "Characterizing the growing microorganisms at species level in 46 anaerobic digesters at Danish wastewater treatment plants: A six-year survey on microbiome structure and key drivers": Additonal file 1: Supplementary tables and figures

Figure S18 Partial least squares estimation of key variables with non-growing bacterial species in MAD.

**Table S1. Overview of WWTP digester capacities, type, and industrial load.**

| WWTP | WWTP capacity (PE)[1] | Number of digesters | Total volume ($m^3$) | AD type | WWTP industrial load | PS/AS in feed[2] |
|---|---|---|---|---|---|---|
| Avedøre | 400,000 | 4 | 24,000 | Mesophilic | NA | + / + |
| Damhusaaen | 350,000 | 4 | 7,600 | Mesophilic | 5% | + / + |
| Egaa | 120,000 | 1 | 3,000 | Mesophilic | NA | + / + |
| Ejby Mølle | 410,000 | 4 | 11,200 | Mesophilic | NA | + / + |
| Esbjerg West | 290,000 | 4 | 9,700 | Mesophilic[3] | 60% | + / + |
| Fornæs | 65,000 | 1 | 1,350 | Mesophilic | 8% | - / + |
| Hjørring | 160,000 | 2 | 2,400 | Mesophilic | 30-50% | + / + |
| Hobro | 30,000 | 1 | 1,800 | Mesophilic | NA | + / + |
| Mariagerfjord | 60,000 | 1 | 2,000 | Mesophilic | NA | - / + |
| Mølleaaværket | 135,000 | 5 | 5,000 | Mesophilic | NA | + / + |
| Randers | 130,000 | 2 | 4,800 | Mesophilic | NA | + / + |
| Ringkøbing | 42,000 | 1 | 880 | Mesophilic | NA | + / + |
| Slagelse | 115,000 | 2 | 3,000 | Mesophilic | 30-50% | + / + |
| Søholt | 99,600 | 1 | 2,100 | Mesophilic | 35% | - / + |
| Viborg | 80,000 | 2 | 3,200 | Mesophilic | 10% | + / + |
| Fredericia | 420,000 | 2 | 4,000 | THP-Mesophilic | 63% | - / + |
| Næstved | 65,000 | 1 | 1,600 | THP-Mesophilic | NA | - / + |
| Aaby | 93,000 | 1 | 1,300 | Thermophilic[4] | 20-40% | - / + |
| Aalborg East | 100,000 | 1 | 1,500 | Thermophilic | 10% | + / + |
| Aalborg West | 330,000 | 2 | 5,000 | Thermophilic | 30% | - / + |
| Bjergmarken | 125,000 | 2 | 2,000 | Thermophilic | 20% | - / + |
| Herning | 175,000 | 1 | 2,500 | Thermophilic | NA | + / + |

NA: not available.

[1] PE = Population equivalents.

[2] "+" and "-" indicates yes and no, respectively. PS = Primary sludge, AS = Activated sludge.

[3] Except Esbjerg West, other plants all have continuously stirred tank reactors (CSTR). Esbjerg West configuration consists of a 38°C methanogenic CSTR, followed by a 75°C hydrolysis tank and second step 42°C methanogenic CSTR. The digester biomass was taken from the first step CSTR.

[4] Aaby operated under thermophilic conditions during Jan. 2013 to Jan. 2016.

**Table S2. Linear regression of key variables individually against alpha diversity using the Shannon diversity index at MAD.**

| Category | Variable | Bacteria | | | | | | Archaea | | |
|---|---|---|---|---|---|---|---|---|---|---|
| | | Total bacteria | | | Growing bacteria | | | | | |
| | | $R^2$ | FDR *P* | Signif. codes | $R^2$ | FDR *P* | Signif. codes | $R^2$ | FDR *P* | Signif. codes |
| Operational variables | Temperature | 0.25 | < 0.001 | *** | 0.16 | < 0.001 | *** | 0.01 | 1 | NS |
| | OLR | 0.09 | 0.24 | NS | 0.31 | < 0.001 | *** | -0.02 | 1 | NS |
| | SRT | 0.05 | 0.10 | NS | 0.06 | 0.04 | * | 0.04 | 0.78 | NS |
| Performance parameters | pH | 0.18 | < 0.001 | *** | 0.15 | < 0.001 | *** | 0.00 | 1 | NS |
| | TAN | 0.07 | 0.003 | ** | 0.15 | < 0.001 | *** | 0.00 | 1 | NS |
| | Alkalinity | 0.02 | 0.17 | NS | 0.08 | < 0.001 | *** | 0.09 | P < 0.001 | *** |
| | TS | 0.005 | 1 | NS | 0.02 | 0.26 | NS | 0.00 | 1 | NS |
| | VS | 0.01 | 0.43 | NS | 0.03 | 0.03 | * | 0.00 | 1 | NS |
| | Total VFA | 0.02 | 1 | NS | 0.02 | 1 | NS | -0.03 | 1 | NS |
| | Acetate | 0.01 | 1 | NS | 0.01 | 1 | NS | -0.02 | 1 | NS |
| | Biogas yield | 0.15 | 0.09 | NS | 0.46 | < 0.001 | *** | -0.04 | 1 | NS |
| | Methane content | -0.01 | 1 | NS | -0.01 | 0.64 | NS | 0.01 | 1 | NS |

OLR: Organic loading rate; SRT: Solids retention time; TAN: Total ammonia nitrogen; TS: Total solids); VS: Volatile solids; Total VFA: Volatile fatty acid; MAD = Mesophilic AD; FDR: False discovery rate. *: FDR *P* < 0.05; **: FDR *P* < 0.01; ***: FDR *P* < 0.001; NS: Not significant.

**Table S3. Permutational multivariate analysis (using continuous variables only) of variance of beta diversity using weighted UniFrac matrix at MAD.**

| Category | Variable | Bacteria | | | | | | Archaea | | |
|---|---|---|---|---|---|---|---|---|---|---|
| | | Total bacteria | | | Growing bacteria | | | | | |
| | | $R^2$ | FDR *P* | Signif. codes | $R^2$ | FDR *P* | Signif. codes | $R^2$ | FDR *P* | Signif. codes |
| Operational variables | Temperature | 0.12 | 0.001 | *** | 0.14 | 0.001 | *** | 0.13 | 0.004 | ** |
| | OLR | 0.21 | 0.001 | *** | 0.21 | 0.001 | *** | 0.13 | 0.029 | * |
| | SRT | 0.04 | 0.001 | *** | 0.04 | 0.001 | *** | 0.03 | 0.17 | NS |
| Performance parameters | pH | 0.04 | 0.001 | *** | 0.05 | 0.001 | *** | 0.04 | 0.016 | * |
| | TAN | 0.13 | 0.001 | *** | 0.17 | 0.001 | *** | 0.07 | 0.004 | ** |
| | Alkalinity | 0.05 | 0.001 | *** | 0.06 | 0.001 | *** | 0.03 | 0.019 | * |
| | TS | 0.04 | 0.001 | *** | 0.05 | 0.001 | *** | 0.02 | 0.029 | * |
| | VS | 0.02 | 0.001 | *** | 0.03 | 0.001 | *** | 0.05 | 0.004 | ** |
| | Total VFA | 0.04 | 0.014 | ** | 0.04 | 0.025 | * | 0.05 | 0.12 | NS |
| | Acetate | 0.03 | 0.031 | * | 0.03 | 0.04 | * | 0.18 | 0.038 | * |
| | Biogas yield | 0.34 | 0.001 | *** | 0.31 | 0.001 | *** | 0.23 | 0.008 | ** |
| | Methane content | 0.08 | 0.04 | * | 0.03 | 0.037 | * | 0.13 | 0.009 | ** |

OLR: Organic loading rate; SRT: Solids retention time; TAN: Total ammonia nitrogen; TS: Total solids; VS: Volatile solids; Total VFA: Volatile fatty acid; MAD = Mesophilic AD; FDR: False discovery rate. *: FDR *P* < 0.5; **: FDR *P* < 0.1; ***: FDR *P* < 0.01; NS: Not significant.

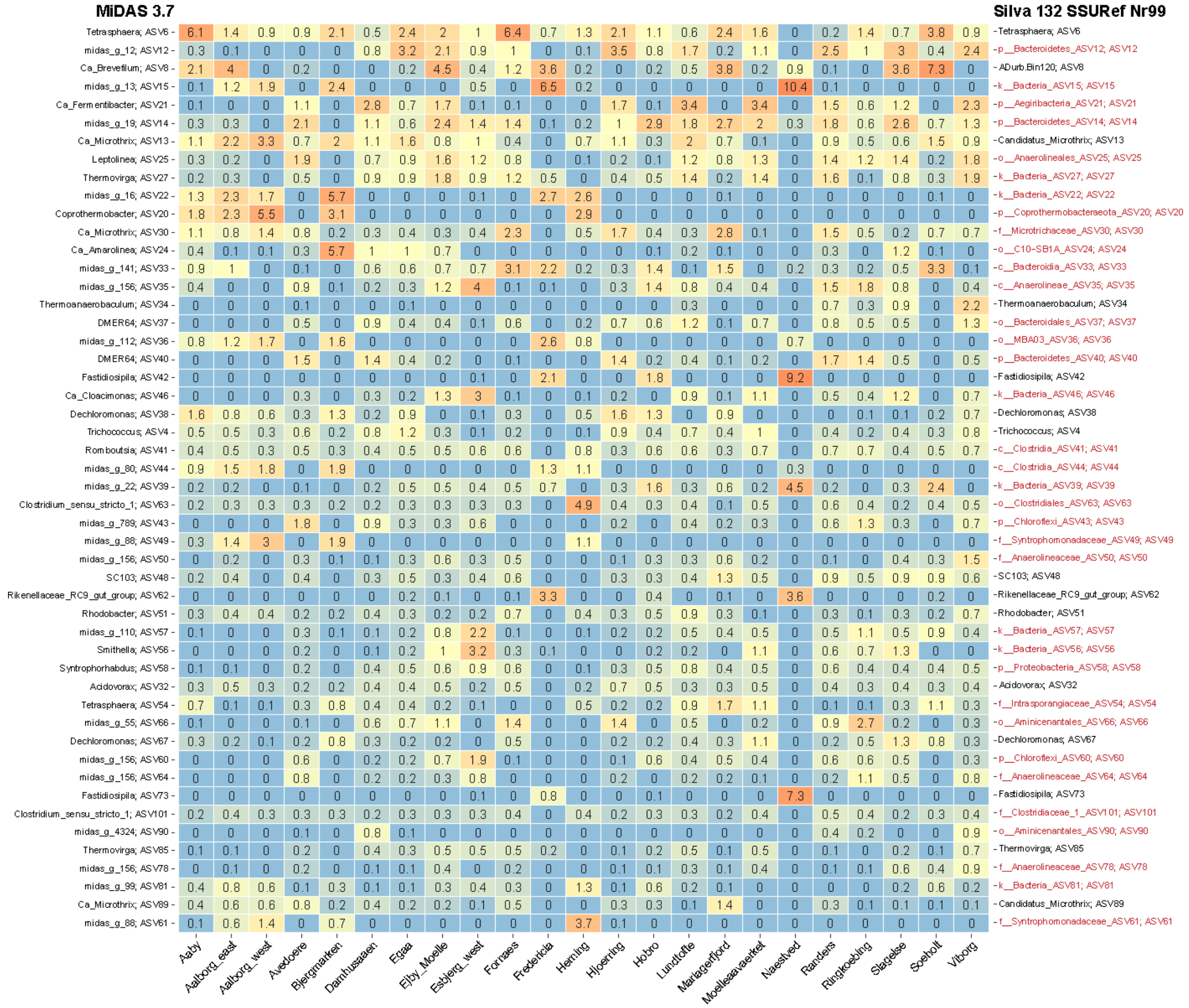


**Figure S1 Comparison of classification between full-length amplicon sequence variants (FL-ASVs) database and the SILVA_132_SSURef_Nr99 database on the top 50 ASVs in the digester sludge samples from Danish wastewater treatment plants.** ASVs not classified at the genus level have been highlighted in red.

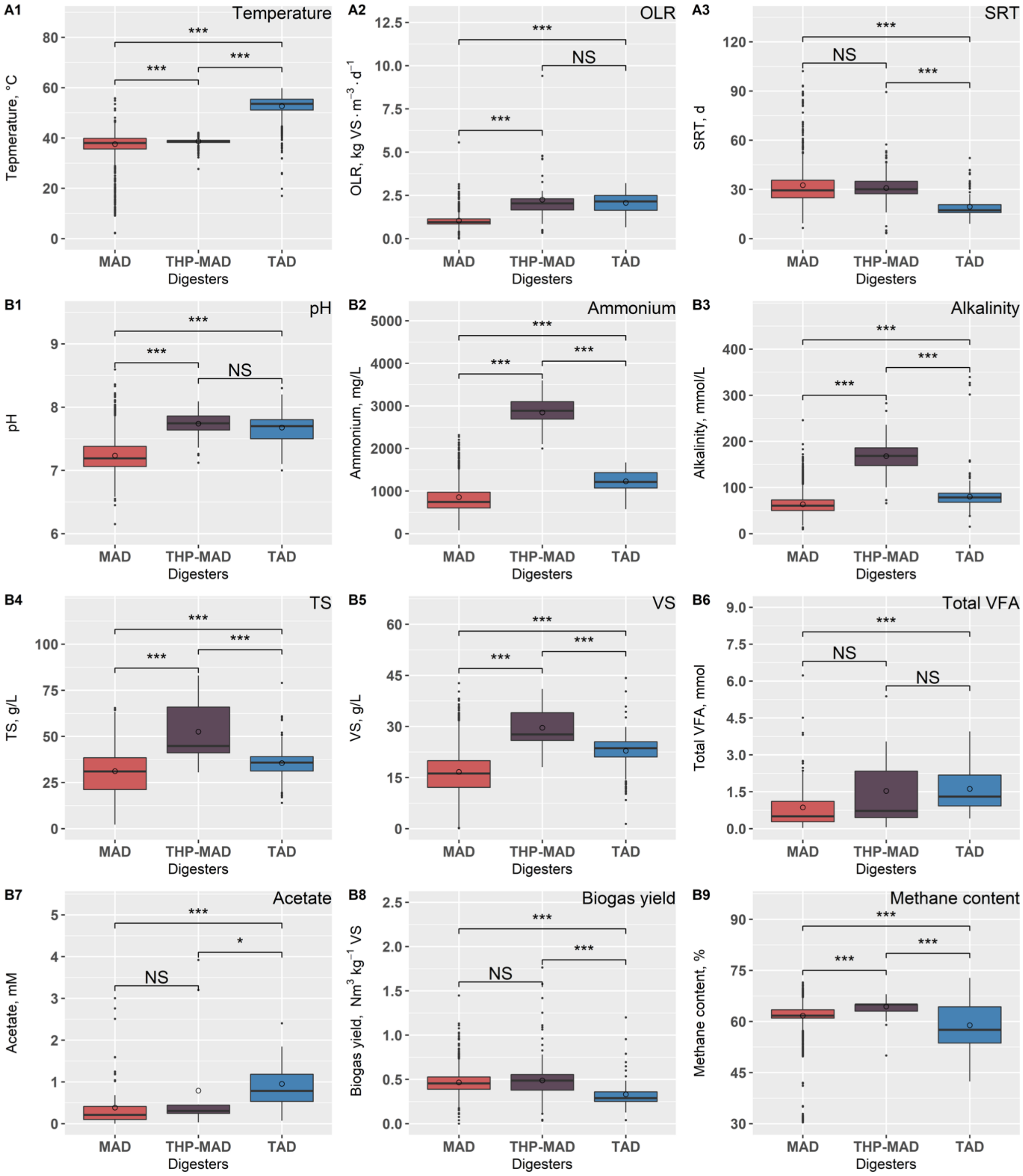


**Figure S2 Box plots of operational and performance parameters of three types of AD.** (A1-A3) Operational parameters: temperature, OLR (organic loading rate), and SRT (solids retention time) (B1-B9). Performance parameters: pH, TAN (total ammonia nitrogen), alkalinity, TS (total solids), VS (volatile solids), Total VFA (volatile fatty acid), acetate, biogas yield, and methane content. MAD = Mesophilic AD, THP-MAD = Mesophilic with thermal hydrolysis pretreatment, TAD = Thermophilic AD. Significant differences are indicated (Dunn test with Bonferroni p value; *, $p < 0.05$; **, $p < 0.01$; ***, $p < 0.001$; NS = Not significant).

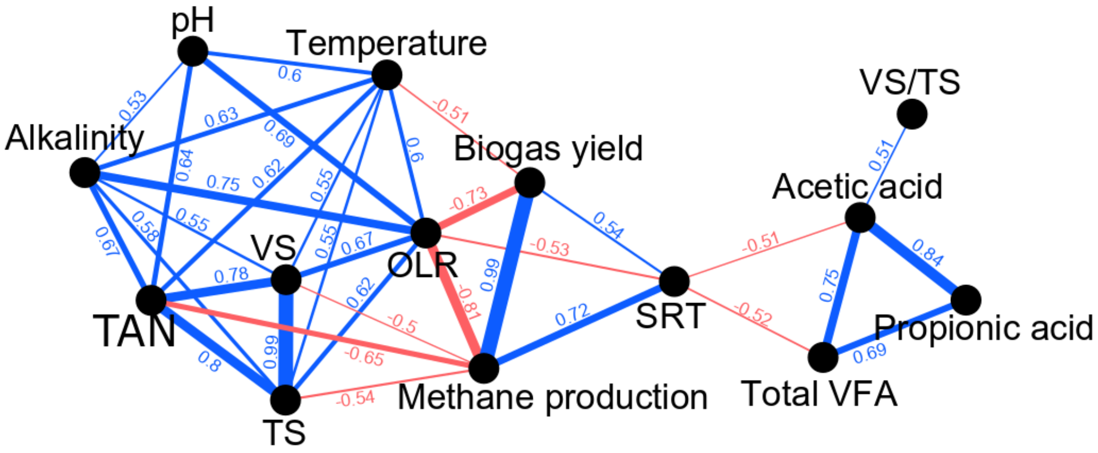


**Figure S3 Spearman correlations on operational and performance parameters in AD.** Positive correlations are in blue, negative correlations are in red. The numbers indicate the coefficients, FDR $P < 0.05$. TAN = Total ammonia nitrogen, OLR = organic loading rate, SRT = solids retention time, TS = total solids, VS = volatile solids, Total VFA = total volatile fatty acid; FDR: False discovery rate.

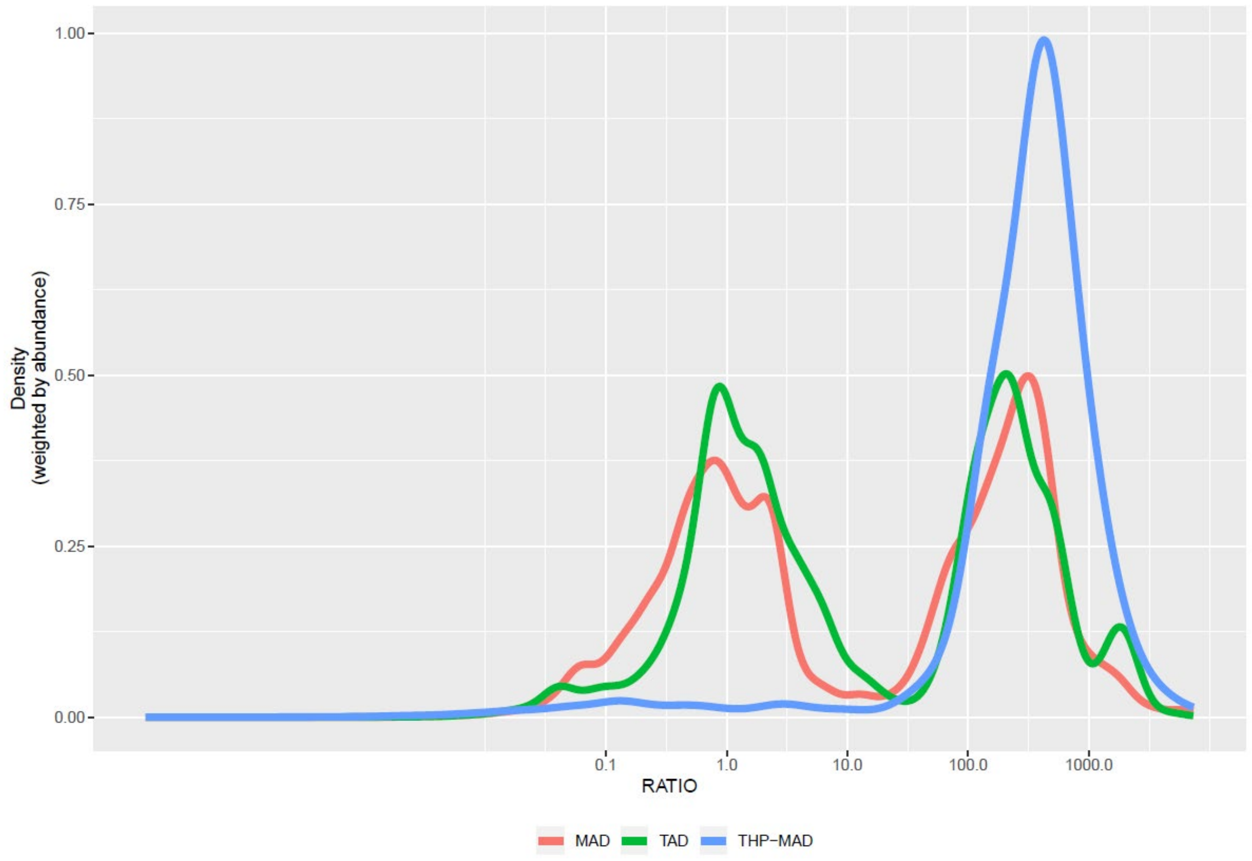


**Figure S4 Distribution of digester:feed relative read abundance ratios for each ASV.** The group with ratio >10 represents the ASVs likely enriched in the digester, compared to the feed sludge, and named as growing group. The read abundance of the ASVs in the group with ratio <10 is unchanged or lower compared to the feed sludge, thus named as non-growing. MAD = Mesophilic AD, THP-MAD = Mesophilic with thermal hydrolysis pretreatment, TAD = Thermophilic AD.

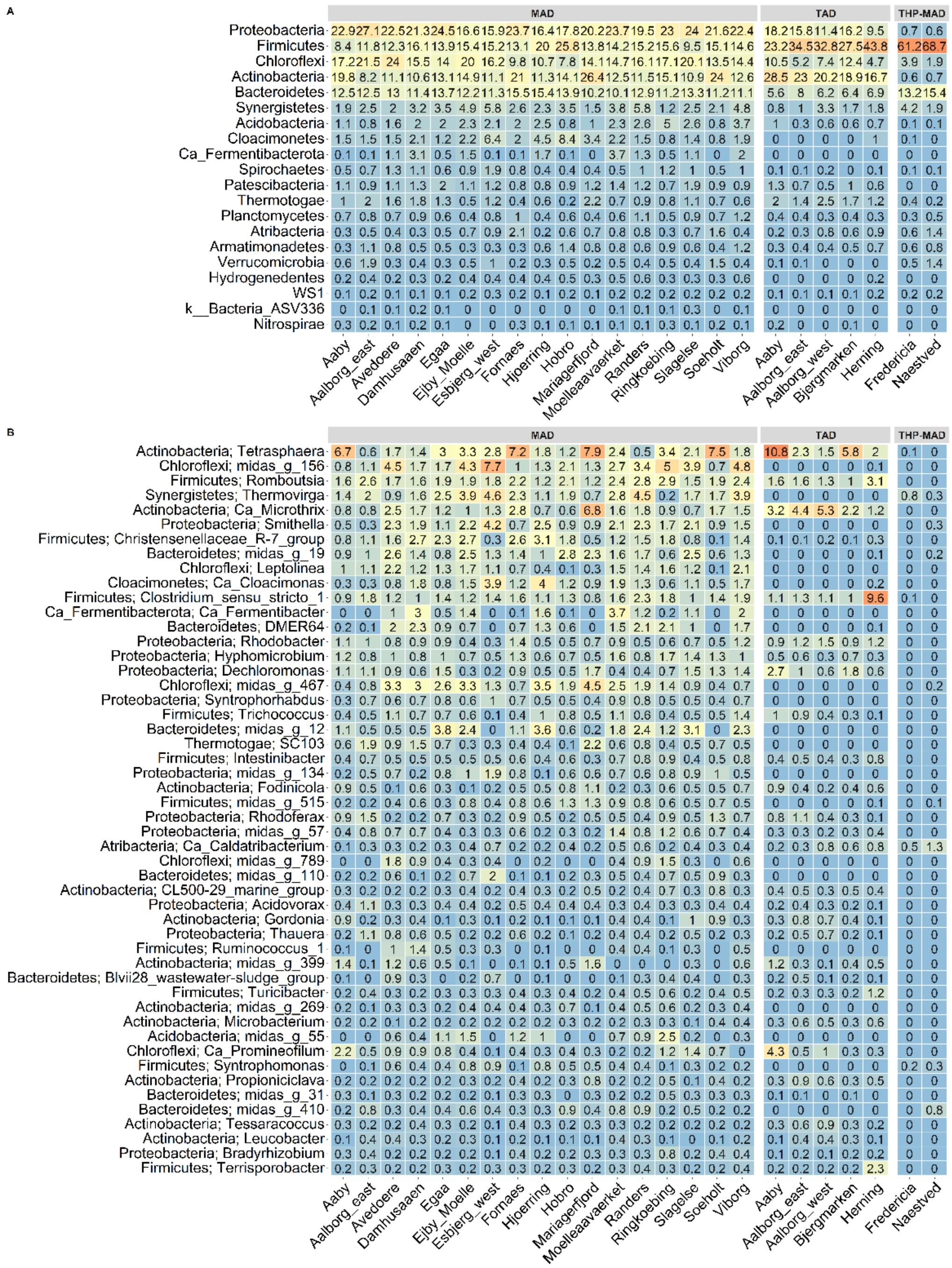


**Figure S5 (A) Relative abundance of the 20 most abundant bacterial phyla in AD. (B) Relative abundance of the 50 most abundant bacterial genera in AD (n = 564).** The median read abundance is shown for each plant. The taxa are sorted by median read abundance across the plants at the respective phylogenetic level (phylum, genus).

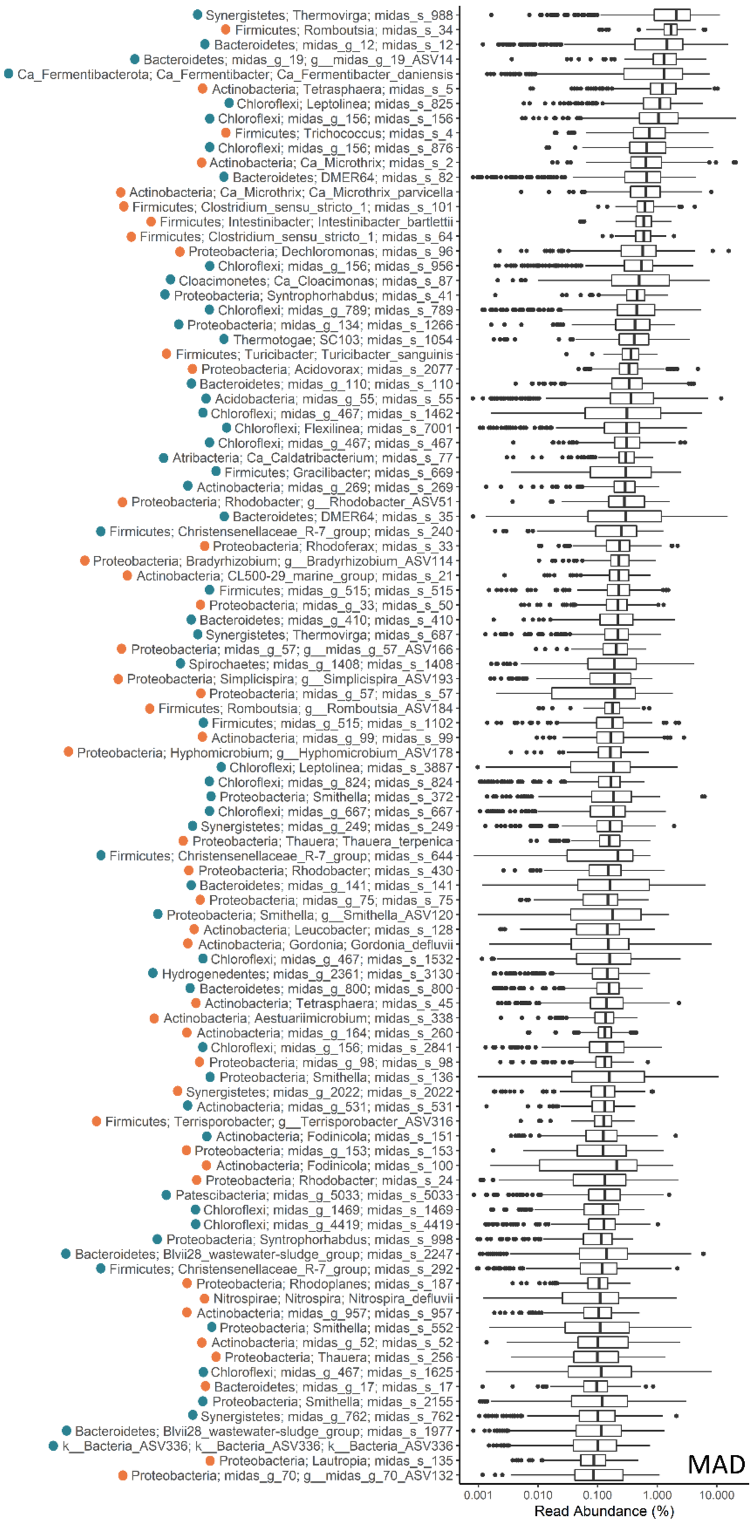


**Figure S6 Boxplots of the top 100 species/ASVs in MAD.** The dots at the left indicate whether the species/ASVs are growing (ratio >10, blue), non-growing or dying off (ratio <10, orange). MAD = Mesophilic AD. Sequences not possessing species-levels classification are shown here as individual ASVs.

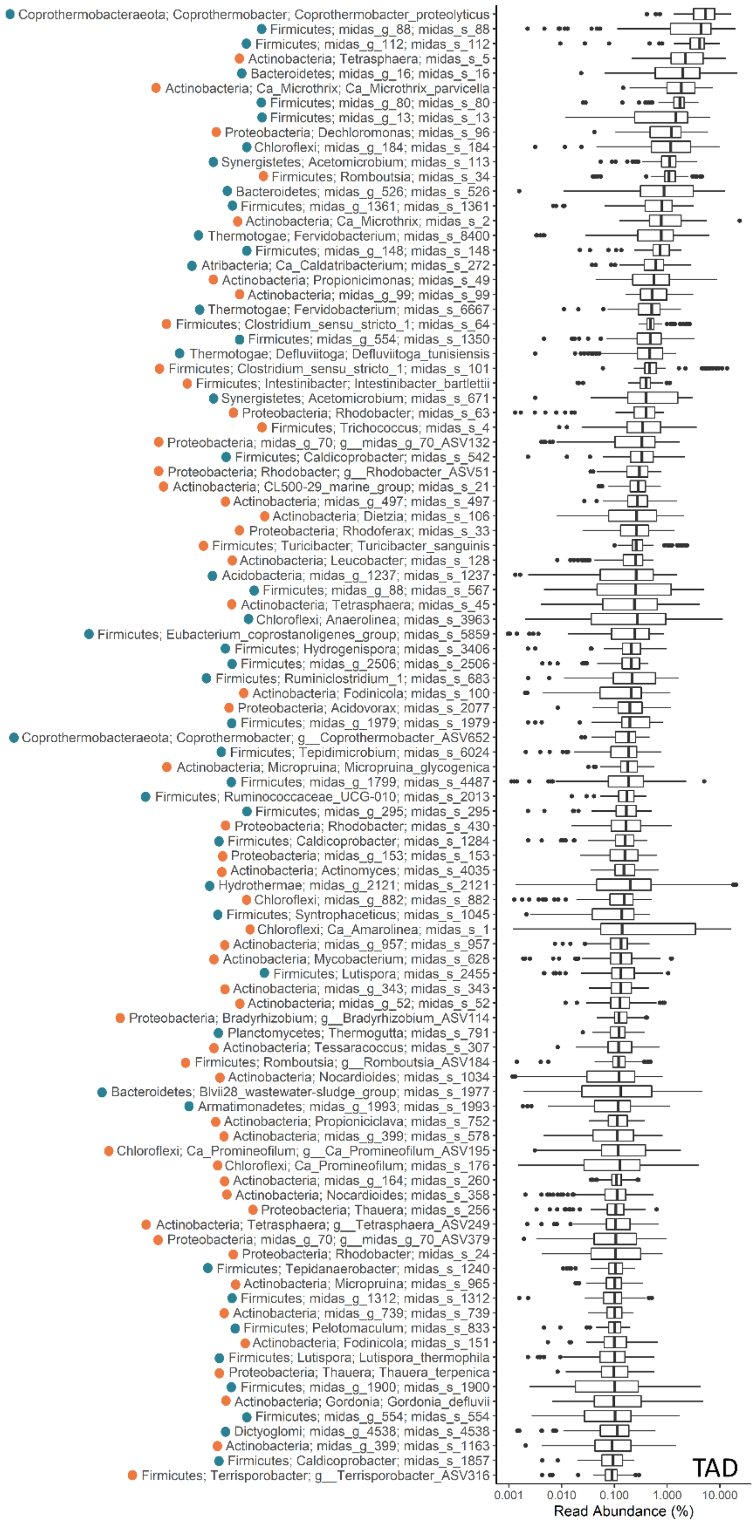


**Figure S7 Boxplots of the top 100 species/ASVs in TAD.** The dots on the left indicate whether the species/ASVs are growing (ratio >10, blue), non-growing or dying off (ratio <10, orange). TAD = Thermophilic AD. Sequences not possessing species-levels classification are shown here as individual ASVs.

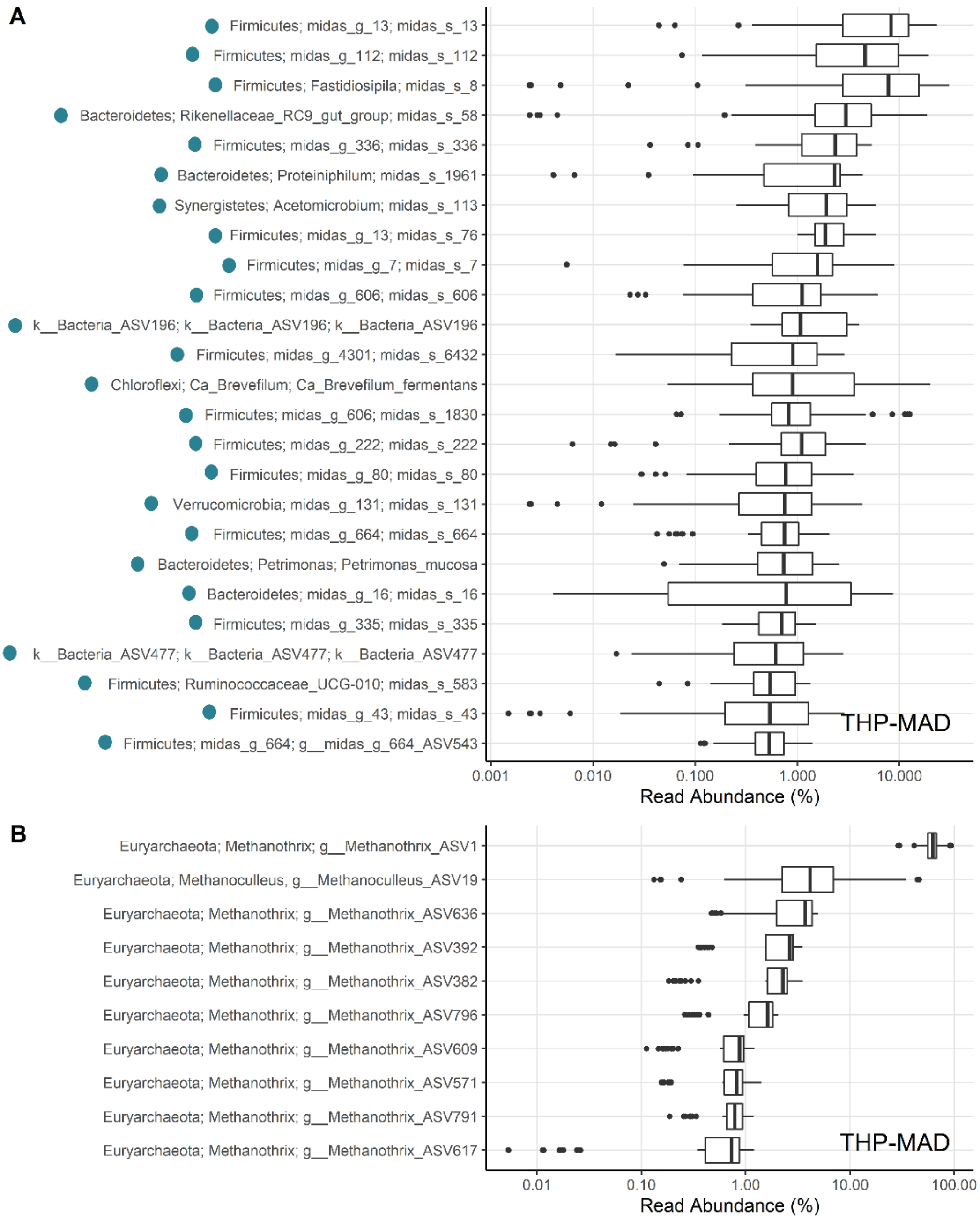


**Figure S8 Boxplots of the most abundant species/ASVs in THP-MAD.** (A) The top 25 most abundant bacterial species/ASVs, (B) The top 10 most abundant archaeal species/ASVs. THP-MAD = Mesophilic with thermal hydrolysis pretreatment. The dots on the left indicate whether the species/ASVs are growing (ratio >10, blue), non-growing or dying off (ratio <10, orange). Sequences not possessing species-levels classification are shown here as individual ASVs.

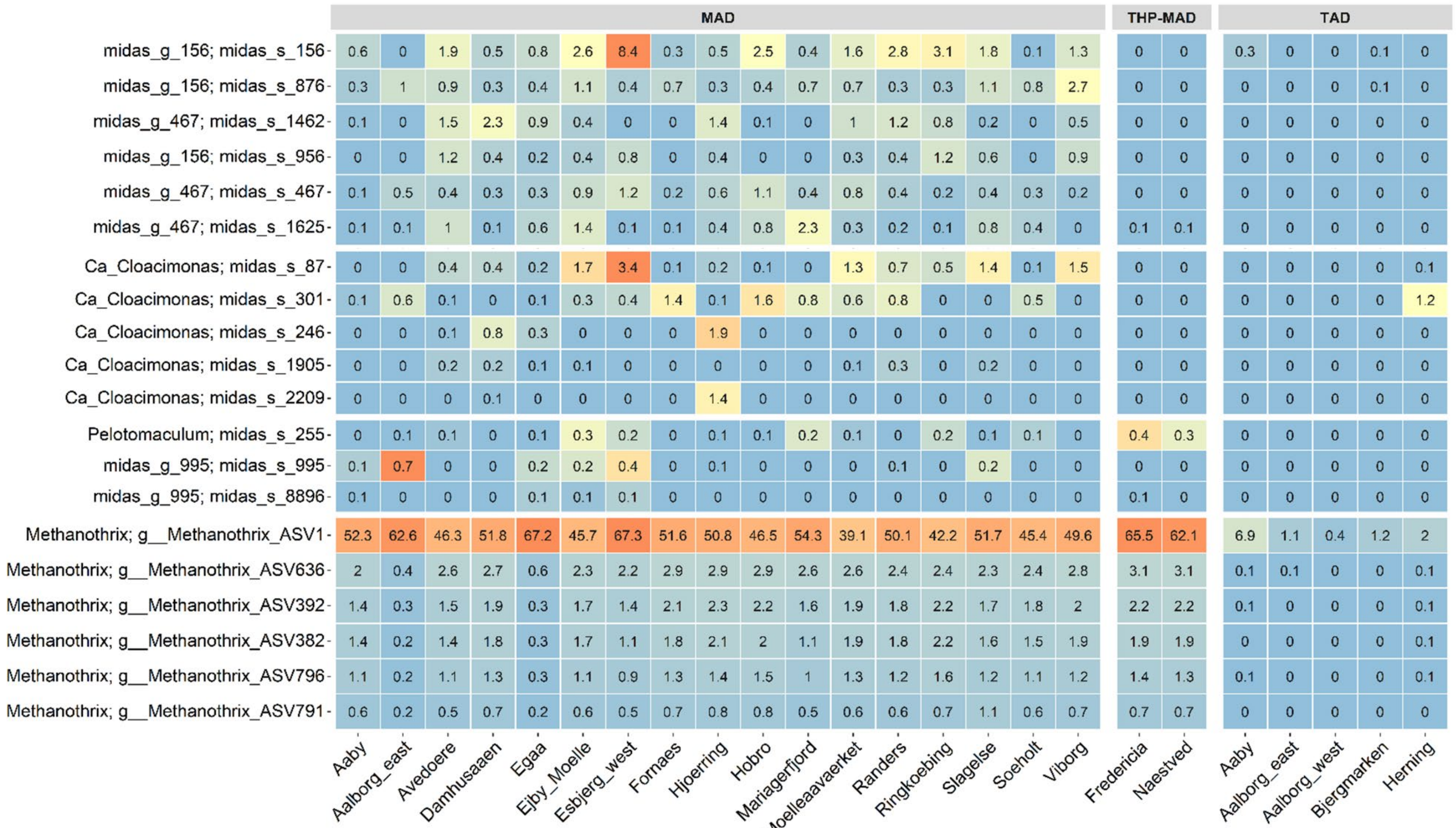


**Figure S9 Heatmap of the most abundant species/ASVs belonging to the genus T78 in MiDAS 2 (split into the genera midas_g_156 and midas_g_467, all family Anaerolineaceae), genus *Ca*. Cloacimonas, genus *Pelotomaculum*, midas_g_995, and genus *Methanothrix* in Danish ADs at WWTPs.** Data represents median values for each plant. MAD = Mesophilic AD, THP-MAD = Mesophilic with thermal hydrolysis pretreatment process, TAD = Thermophilic AD.

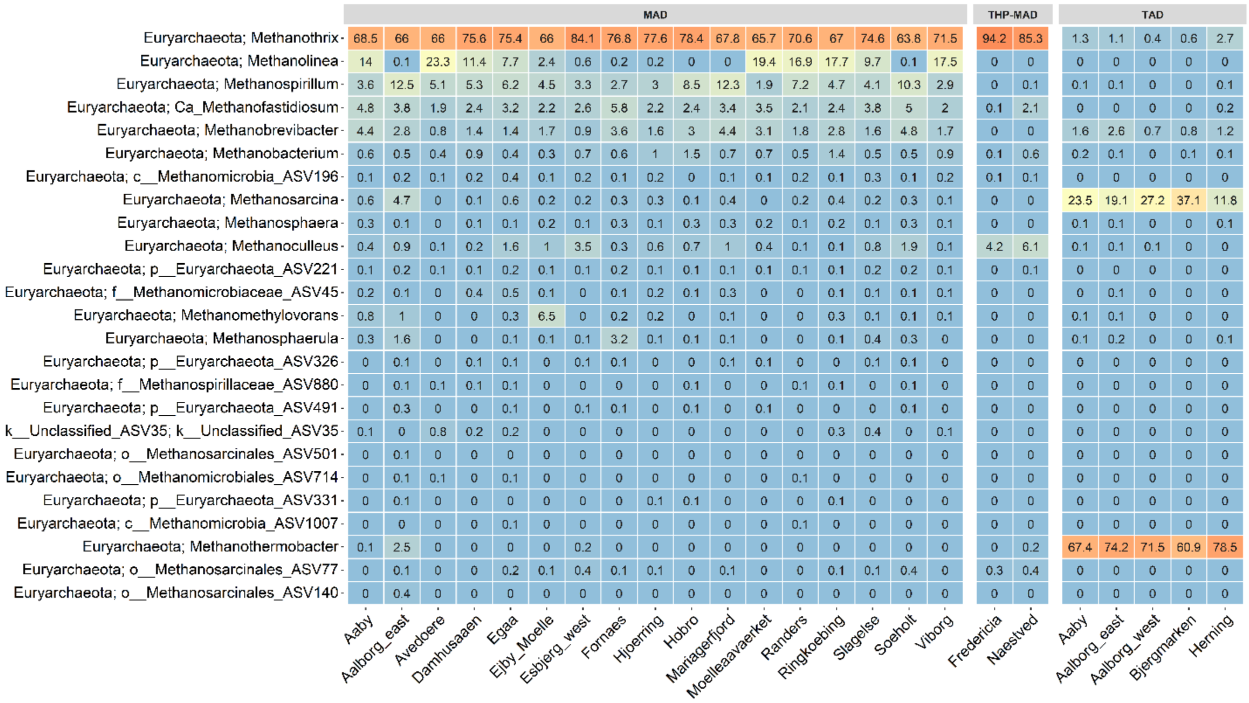


**Figure S10 Relative abundance of the 25 most abundant archaeal genera in AD (n = 402).** The median read abundance is shown for each plant. The taxa are sorted by median read abundance across the plants at the respective phylogenetic level (phylum, genus).

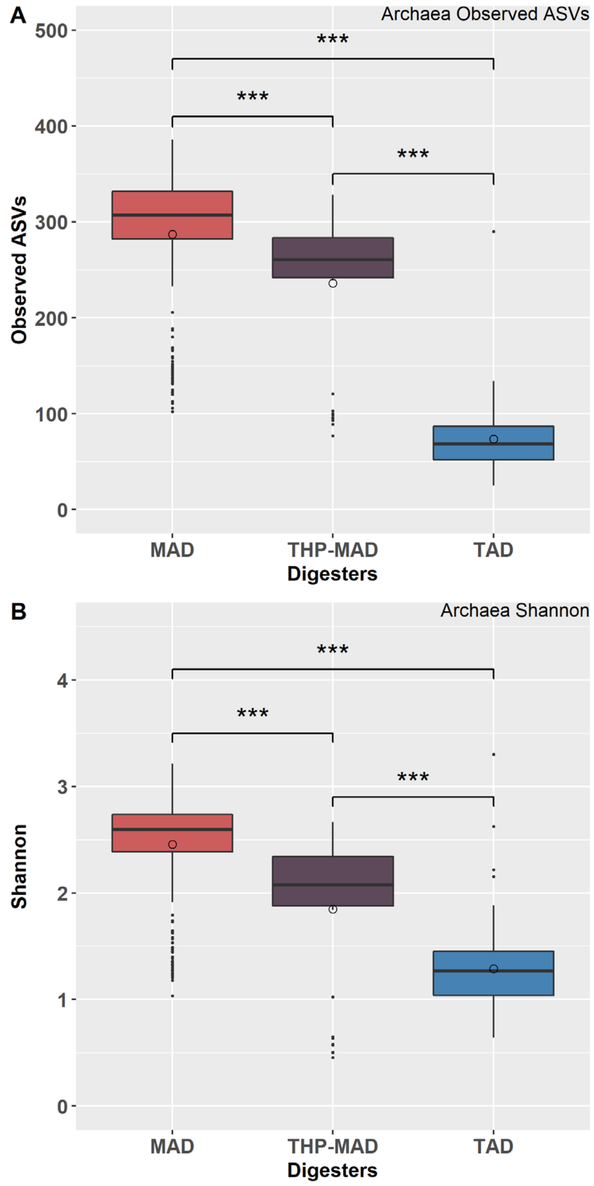


**Figure S11 Boxplots of alpha diversity measures of archaeal community of three types of AD.** (A) Observed ASVs of archaeal community, (B) Shannon index of archaeal community. MAD = Mesophilic AD, TAD = Mesophilic AD, THP = Mesophilic with thermal hydrolysis pretreatment process. Significant differences are indicated (Wilcoxon rank-sum test; *, $p < 0.05$; **, $p < 0.01$; ***, $p < 0.001$).

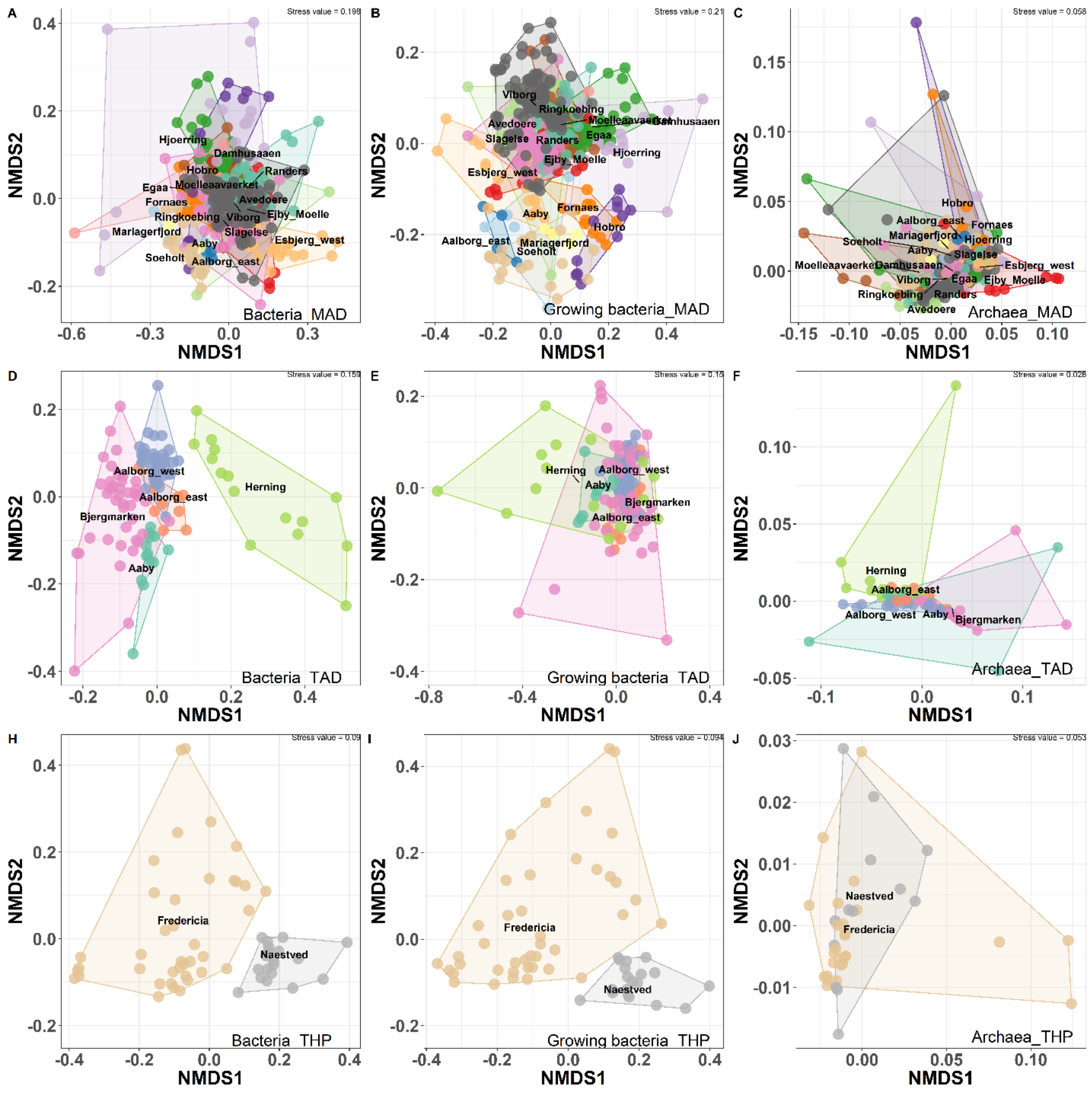


**Figure S12 Non-metric multidimensional scaling (NMDS) plots of bacterial and archaeal community structure based on weighted Unifrac matrix colored by WWTPs.** (A, D, H) The entire bacterial communities colored by WWTPs in MAD, TAD, and THP. (B, E, I) The growing bacterial communities colored by WWTPs in MAD, TAD, and THP. (C, F, J) The archaeal communities colored by WWTPs in MAD, TAD, and THP. MAD = Mesophilic AD, TAD = Mesophilic AD, THP = Mesophilic with thermal hydrolysis pretreatment process.

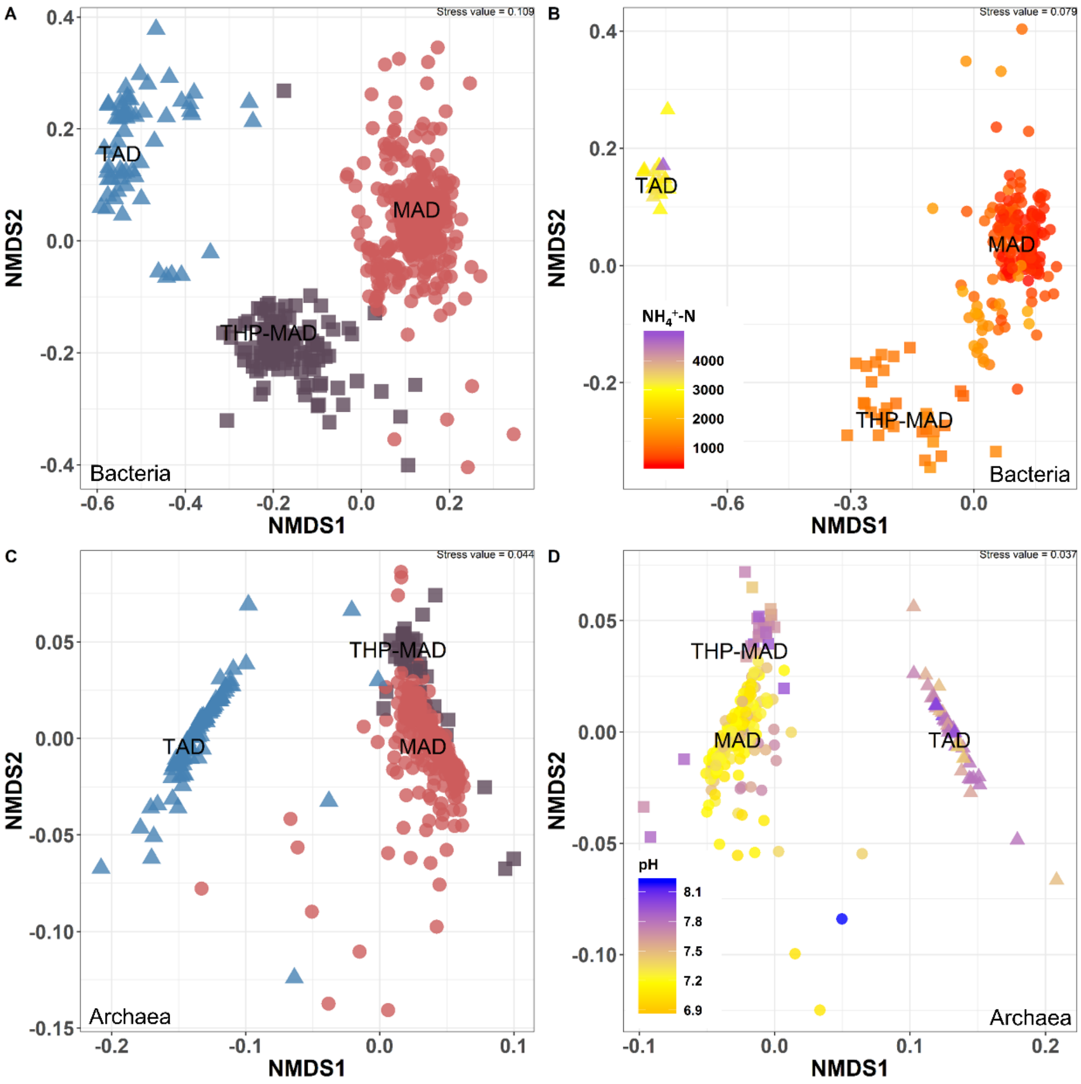


**Figure S13 Non-metric multidimensional scaling (NMDS) plots of all Bacteria and Archaea community structures based on weighted Unifrac matrix.** (A) The bacterial communities colored by type of AD (n = 564), (B) The bacterial communities colored by concentrations of TAN nitrogen (n = 234), (C) The archaeal communities colored by type of AD (n = 402), (D) The archaeal communities colored by pH values (n = 255). MAD = Mesophilic AD, THP-MAD = Mesophilic with thermal hydrolysis pretreatment, TAD = Thermophilic AD, TAN = Total ammonia nitrogen, mg N/L.

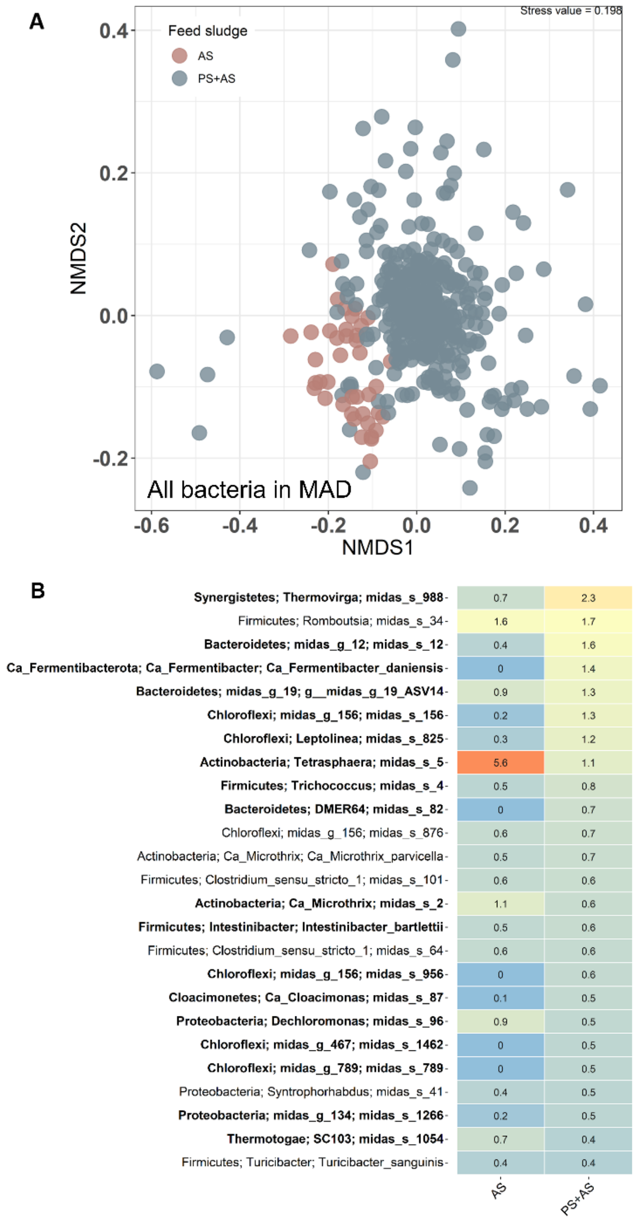


**Figure S14 (A) Non-metric multidimensional scaling (NMDS) plots of entire bacterial community structure based on weighted UniFrac matrix in MAD. (B) Heatmap of 25 most abundant bacterial species in MAD digesters depending on the composition of feed sludge (AS and PS).** The median read abundance is shown for each group. Species in bold indicate significant differences of median relative abundance between the two groups (Wilcoxon rank-sum test, $P < 0.05$). AS = Activated sludge; PS = Primary sludge.

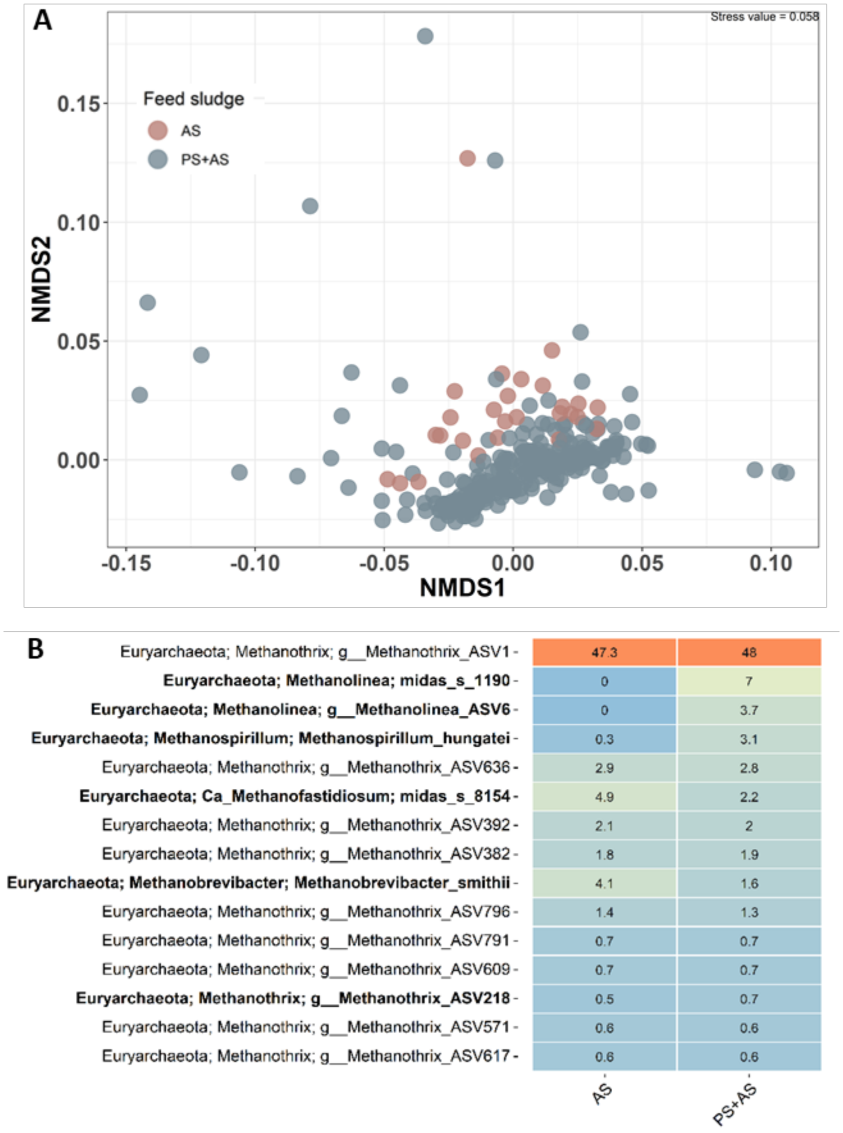


**Figure S15 (A) Non-metric multidimensional scaling (NMDS) plots of archaeal community structure based on weighted UniFrac matrix in MAD. (B) Heatmap of 15 most abundant archaeal species in MAD digesters depending on the composition of feed sludge (AS and PS).** The median read abundance is shown for each group. Species in bold indicate significant differences of median relative abundance between the two groups (Wilcoxon rank-sum test, $P < 0.05$). AS = Activated sludge; PS = Primary sludge.

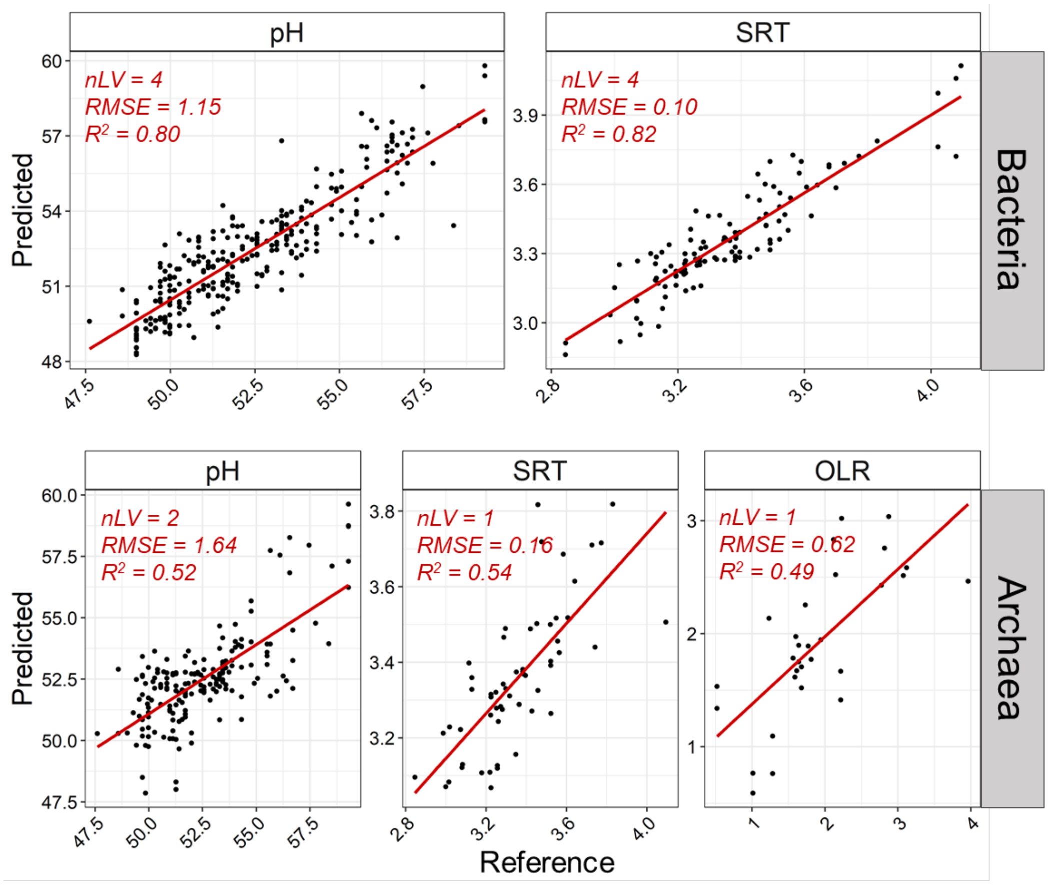


**Figure S16 Prediction plots of other important parameters based on bacterial and archaeal microbiome in MAD by partial least square regression.** MAD = mesophilic AD, OLR = organic loading rate, TAN = total ammonia nitrogen. nLV = number of selected components, RMSE = root mean squared error.

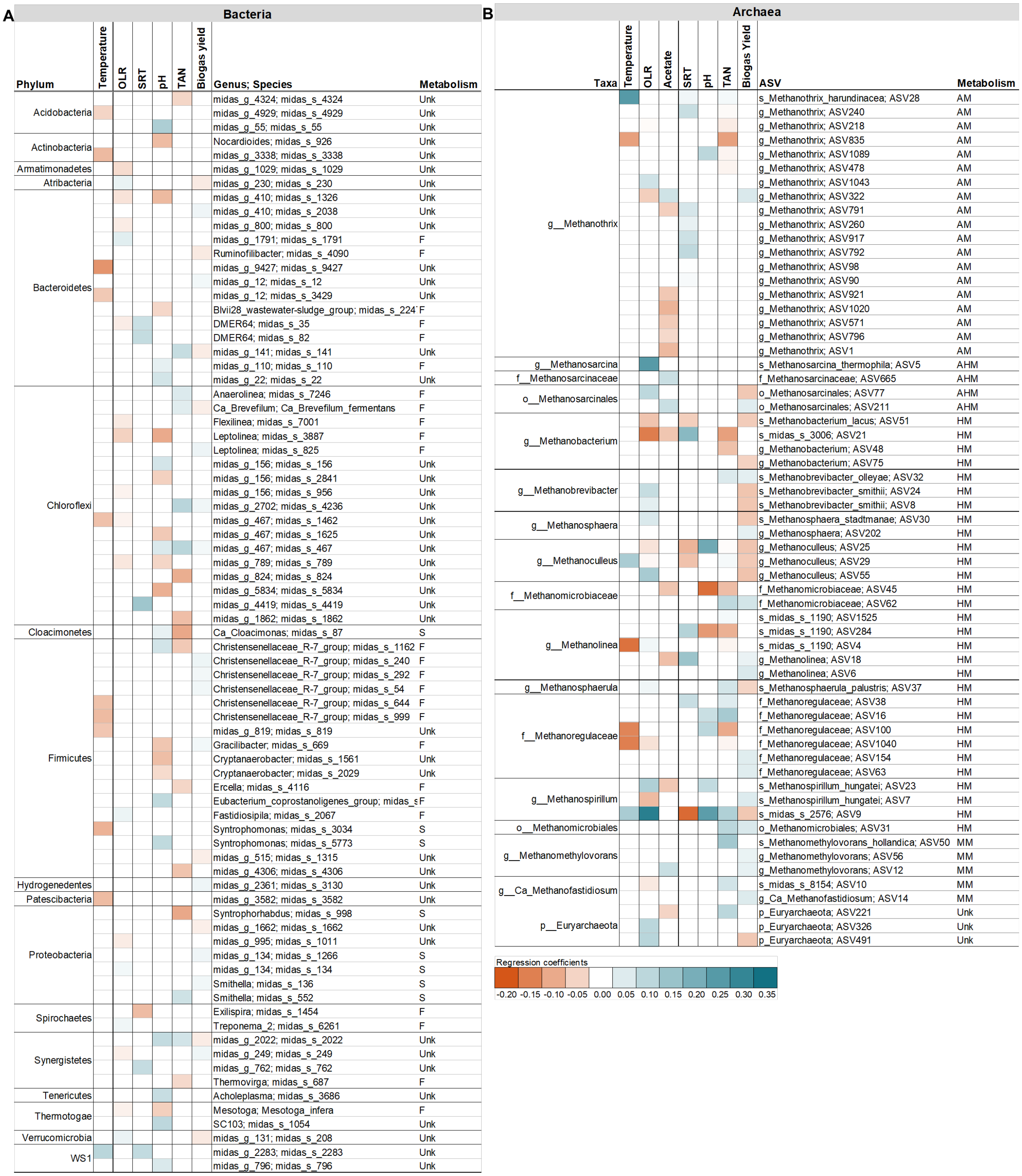


**Figure S17 Complete list for partial least squares estimation of key variables with growing bacterial species (A) and archaeal ASVs (B) in MAD.** P < 0.05, positive correlation in blue, negative correlation in orange.

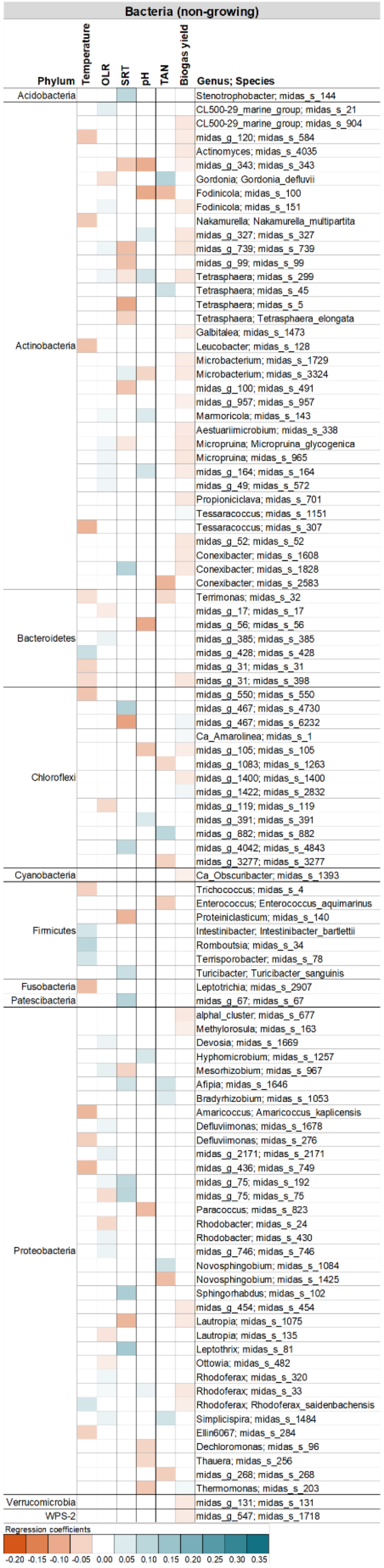


**Figure S18 Partial least squares estimation of key variables with non-growing bacterial species in MAD.** P < 0.05, positive correlation in blue, negative correlation in orange.
