## Supplementary material for "Characterizing the growing microorganisms at species level in 46 anaerobic digesters at Danish wastewater treatment plants: A six-year survey on microbiome structure and key drivers": Additonal file 2: Characterization of key parameters of ADs

**Additional file 2: Characterization of key parameters of anaerobic digestions (ADs)**

### 1. Operational parameters

Main AD operational parameters are temperature, SRT (equivalent to hydraulic retention time (HRT) in CSTRs), and OLR. Temperature is a key parameter in AD as it affects the degradation rates, the physicochemical equilibria as well as the growth rates and metabolism of microorganisms [1]. The median temperature of the three types of digesters were 38.0°C, 38.6°C, and 53.6°C, for MAD, THP-MAD, and TAD, respectively. Anaerobic digesters are sized according to the SRT to ensure that substrate can be efficiently converted to methane. The median SRT of the three types of digester were 32.6, 30.8, and 19.5 days. Other surveys have also shown same trend, where digesters at WWTPs had a wide range of SRT from 9 to 100 days [2–4]. In CSTRs, OLR is linked to the SRT and the substrate concentration. For a fixed substrate concentration, the high SRT implies low OLR. In our survey, MAD generally ran at a relatively lower OLR than THP-MAD and TAD (**Figure S1.A2**), which was within the range of mesophilic full-scale digesters at WWTPs in other European countries (0.70 ~ 1.90 kg VS/m$^3$·d) [2,4–7]. Other THP-mesophilic full-scale digesters at WWTPs are reported to run at an OLR around 3 to 4.2 kg VS/m$^3$·d [8,9], as a consequence of the lower SRT (~ 15 day).

### 2. Performance parameters

The performance parameters (pH, total ammonia nitrogen (TAN), alkalinity, TS, VS, and biogas yield) were found to be significantly different across all three types of AD (**Figure S1**). TAN is a key physicochemical parameter in digesters, which is not only an essential macronutrient for microbial growth but also a common indicator for AD inhibition. Lower TAN concentrations are typically observed in WWTP-based AD than in manure and food waste-based AD [2,4,10–12]. The digesters surveyed showed a median TAN of 745 mg/L, similar to other full-scale mesophilic AD studies at WWTP, where the average concentration was 1026 mg/L [2,4,6,7,10,11,13–15]. TAD showed a median of 1215 mg/L, as consequence of the higher OLR and faster degradation of rates at the thermophilic range [16]. THP-MAD had higher median TAN concentration of 2888 mg/L due to the thermal hydrolysis of waste activated sludge, probably releasing more protein. These TAN concentrations are much lower than the reported inhibitory threshold for TAN ( > 5000 mg N/L) [17,18]. Moreover, the low VFA concentrations in the THP-MAD digesters and biogas yield similar to the MAD suggest that inhibition was not noticeable. The pH of the three

AD types ranged around neutrality (**Table 1**) and were similar to other studies considering mesophilic (6.98 ~ 7.90) [2–7,10,11,13–15] and thermophilic (7.2 ~ 8.00) [3,4,11,15]. The optimal range reported for alkalinity is 100 ~ 300 mM as $CaCO_3$ [19]. The alkalinity concentrations of MAD and TAD were a bit lower than the optimum reported range (**Table 1**), although instability problems were not noticed in any plant during the six-year survey. The higher concentration of alkalinity in THP-MAD is explained by the higher TAN values.

The TS and VS of the digesters are linked to the nature of the substrate and the degradation efficiency. Different ranges are found for different digester configurations, however, the ratio of VS to TS was around 59% and similar among AD on the WWTPs (**Table 1**). Our observations were similar to other mesophilic WWTP studies, where average TS and VS were 30.3 and 18.0 g/L, and the ratio of VS to TS was 60% [2,5–7,10,13]. The median concentrations of total VFAs were low, 0.50 mM (MAD), 0.73 mM (THP-MAD), and 1.30 mM (TAD). These values are consistent with other full-scale digesters operated at low OLR and long SRT [20,21] and far below the stable cut-off value of VFAs (50 ~ 70 mM) [22,23]. The biogas yield was similar between MAD and THP-MAD, and higher than the TAD. However, taking into account the methane content in the biogas, the methane production of three types were 0.27 (MAD), 0.65 (THP-MAD), and 0.36 (TAD) $Nm^3/m^3d$, similar to reported biogas productions in MAD and TAD systems [4,14–16,23].
